# An efficient peptide ligase engineered from a bamboo asparaginyl endopeptidase

**DOI:** 10.1101/2023.09.07.556767

**Authors:** Xin-Bo Wang, Cong-Hui Zhang, Teng Zhang, Hao-Zheng Li, Ya-Li Liu, Zeng-Guang Xu, Gang Lei, Chun-Ju Cai, Zhan-Yun Guo

## Abstract

In recent years, a few asparaginyl endopeptidases (AEPs) from certain higher plants have been identified as efficient peptide ligases with wide applications in protein labeling and cyclic peptide synthesis. Recently, we developed a NanoLuc Binary Technology (NanoBiT)-based peptide ligase activity assay to identify more AEP-type peptide ligases. Herein, we screened 61 bamboo species from 16 genera using this assay and detected AEP-type peptide ligase activity in the crude extract of all tested bamboo leaves. From a popular bamboo species, *Bambusa multiplex*, we identified a full-length AEP-type peptide ligase candidate (BmAEP1) via transcriptomic sequencing. After its zymogen was overexpressed in *Escherichia coli* and self-activated *in vitro*, BmAEP1 displayed high peptide ligase activity, but with considerable hydrolysis activity. After site-directed mutagenesis of its ligase activity determinants, the mutant zymogen of [G238V]BmAEP1 was normally overexpressed in *E. coli*, but failed to activate itself. To solve this problem, we developed a novel protease-assisted activation approach in which trypsin was used to cleave the mutant zymogen and was then conveniently removed via an ion-exchange chromatography. After the non-covalently bound cap domain was dissociated from the catalytic core domain under acidic conditions, the recombinant [G238V]BmAEP1 displayed high peptide ligase activity with much lower hydrolysis activity, and could efficiently catalyze inter-molecular protein ligation and intra-molecular peptide cyclization. Thus, the engineered bamboo-derived peptide ligase represents a novel tool for protein labeling and cyclic peptide synthesis.

## 1. Introduction

Asparaginyl endopeptidases (AEPs, also known as legumains, EC 3.4.22.34) are a group of cysteine proteases widely present in animals and plants [1−4]. In general, AEPs specifically hydrolyze the peptide bond at the C-terminus of Asn or Asp residues. However, in recent years it was discovered that a few AEPs from certain higher plants have evolved peptide ligase activity [4−6], such as butelase-1 from *Clitoria ternatea* (butterfly pea) [7] and OaAEP1 from *Oldenlandia affinis* [8]. The AEP-type peptide ligases (also known as peptidyl asparaginyl ligases, PALs) can efficiently catalyze intra-or inter-molecular peptide ligation via transpeptidation [4−6]. They are responsible for the biosynthesis of certain cyclic peptides from linear precursors [9−11], and have wide applications in biomedical studies [12−15], such as protein and peptide ligation [16−18], labeling [19−23], immobilization [24−26], and cyclization [27−30], as well as in living cell labeling [31−33]. To date, AEP-type peptide ligases have only been identified from a few higher plants, such as *Clitoria ternatea* (butterfly pea) [7], *Oldenlandia affinis* [8], *Helianthus annuus* [34], *Petunia x hybrida* [35], *Viola yedoensis* [36], and *Momordica cochinchinensis* [37,38].

To identify more AEP-type peptide ligases, we recently developed a NanoLuc Binary Technology (NanoBiT)-based peptide ligase activity assay via the introduction of a peptide ligase recognition motif to the C-terminus of an inactive large NanoLuc fragment (LgBiT) and the introduction of a peptide ligase-preferred nucleophilic motif to the N-terminus of a synthetic low-affinity SmBiT complementation tag [39]. In the present study, we screened 61 bamboo species from 16 genera using this activity assay, and detected peptide ligase activity in the crude extract of all tested bamboo leaves. From a popular horticultural bamboo species, *Bambusa multiplex*, we identified a bamboo AEP (designated as BmAEP1) that displayed high peptide ligase activity, but with considerable hydrolysis activity after its zymogen was overexpressed in *Escherichia coli* and self-activated *in vitro*. To eliminate its hydrolysis activity, we conducted site-directed mutagenesis at its ligase activity determinants (LADs), and developed a novel protease-assisted activation approach for *in vitro* activation of the mutant [G238V]BmAEP1 zymogen, which failed to activate itself after bacterial overexpression. The engineered [G238V]BmAEP1 displayed high peptide ligase activity with much lower hydrolysis activity, and could efficiently catalyze inter-molecular protein ligation and intra-molecular peptide cyclization, representing a novel tool for protein labeling and cyclic peptide synthesis.

## 2. Materials and methods

### 2.1. Peptide synthesis

The ligation version SmBiT (GI-SmBiT), neuropeptide substance P (GI-SP), and sunflower trypsin inhibitor (SFTI-NHV) were chemically synthesized at GL Biochem (Shanghai, China) via standard solid-phase peptide synthesis. The crude peptides were purified to homogeneity by high performance liquid chromatography (HPLC) using a C_18_ reverse-phase column (Zorbax 300SB-C18, 9.4 × 250 mm; Agilent Technologies, Santa Clara, CA, USA) and confirmed by mass spectrometry.

### 2.2. Overexpression and purification of the ligation version LgBiT and NanoLuc

The expression construct for the ligation version LgBiT (LgBiT-NHV) was generated in our previous study [39]. The expression constructs for the ligation version NanoLucs (NanoLuc-NHV and NanoLuc-NAL) were generated by site-directed mutagenesis via a QuikChange approach based on our previous construct pET/6×His-NanoLuc-Cys [40], and their nucleotide and amino acid sequences are shown in Fig. S1. After overexpression in the *E. coli* strain BL21(DE3) as soluble proteins, the ligation version LgBiT and NanoLucs were purified by an immobilized metal ion affinity chromatography (Ni^2+^ column) and an ion-exchange chromatography (DEAE column), analyzed by sodium dodecyl sulfate-polyacrylamide gel electrophoresis (SDS-PAGE) and bioluminescence measurement, and stored at −80 °C according to our previous procedures [39,40].

### 2.3. Bioluminescence measurement

NanoLuc substrate stock solution was purchased from Promega (Madison, WI, USA) and stored at −80 °C. Before use, the substrate stock was diluted 100-fold using NanoBiT diluting solution [phosphate-buffered saline (PBS) plus 0.1% bovine serum albumin (BSA) and 0.01% Tween-20]. For bioluminescence measurement, diluted samples were transferred to a 384-well white plate (10 μl/well), mixed with equal volume of the diluted NanoLuc substrate, and immediately measured on a SpectraMax iD3 plate reader (Molecular Devices, Sunnyvale, CA, USA). The measured bioluminescence data are expressed as mean ± standard deviation (SD, *n* = 3) if not mentioned otherwise, and plotted using SigmaPlot10.0 software. The accumulated ligation product was calculated from the measured bioluminescence data according to the measured specific luciferase activity of the ligation version LgBiT after complemented with the high-affinity HiBiT tag.

### 2.4. NanoBiT-based peptide ligase activity assays

The NanoBiT-based peptide ligase activity assays were conducted according to our previous procedure [39]. Briefly, the pre-warmed crude or purified peptide ligase was mixed with the recombinant LgBiT-NHV and the synthetic GI-SmBiT at the indicated concentrations. After incubation at the indicated temperature for the indicated time, 10 μl of the reaction mixture was removed and diluted indicated folds using the NanoBiT diluting solution for subsequent bioluminescence measurement.

### 2.5. Collection of bamboo leaves and screening of peptide ligase activity

Fresh leaves of *Bambusa multiplex* were collected from the campus of Tongji University (Shanghai, China), and other fresh bamboo leaves were collected from Anhui Taiping Experimental Station of International Center for Bamboo and Rattan (Taiping County, Anhui Province, China). To prepare the crude extract, 1.0 g of the fresh bamboo leaves was cut to small pieces and ground in a mortar with 4.0 ml of cold extraction buffer [PBS plus 1.0 mM ethylene glycol tetraacetic acid (EGTA) and 5.0 mM β-mercaptoethanol]. After centrifugation (10000 *g*, 5 min), the supernatant was used for peptide ligase activity assay according to our previous procedure [39].

Briefly, the pre-warmed crude extract (typically 50 μl) was mixed with equal volume of the pre-warmed substrate mixture (10 μM LgBiT-NHV and 20 μM GI-SmBiT). After incubation at 20 °C for the indicated time, 10 μl of the ligation mixture was removed and diluted 2,000-fold using the NanoBiT diluting solution for subsequent bioluminescence measurement.

### 2.6. Transcriptomic sequencing and cDNA cloning

Transcriptomic sequencing of the fresh leaves of *Bambusa multiplex* and subsequent data assembly and analysis were conducted at Biomarker Technologies (Beijing, China). The raw sequence reads have been submitted to the SRA database at NCBI with an accession number of <u>SRX19544757</u>. An AEP-type peptide ligase candidate (BmAEP1) was identified from the deduced proteins according to its homology with butelase-1, and its nucleotide and amino acid sequences are shown in Fig. S2. Signal peptide of the BmAEP1 pre-zymogen was predicted by the online SignalP6.0 algorithm (https://services.healthtech.dtu.dk/service.php?SignalP-6.0). Structure of the BmAEP1 zymogen was predicted by the online RoseTTAFold algorithm (https://robetta.bakerlab.org).

To clone cDNA of the BmAEP1 pre-zymogen, total RNA was isolated from the fresh bamboo leaves using Trizol reagent (Beyotime Biotechnology, Shanghai, China). After reverse transcription via a first-strand cDNA synthesis kit (Beyotime Biotechnology), the reaction mixture was used as the template for polymerase chain reaction (PCR). After thermal cycling using pfu DNA polymerase and two oligo primers (BmAEP1-F1: 5’-ATG GCG TCT ATC CGC CTT CTT CCC-3’; BmAEP1-R1: 5’-TCA AGC ACT AAA ACC CTT GTG GAT GGA-3’), Taq DNA polymerase was added and continually incubated at 72 °C for 30 min. Thereafter, the mixture was subjected to agarose gel electrophoresis, and an ∼1.5 kb DNA fragment was recovered from the gel and ligated into a pUCm-T vector (Beyotime Biotechnology). Finally, the ligation mixture was transformed into *E. coli* strain DH5α and three colonies were subjected to plasmid extraction and DNA sequencing.

### 2.7. Overexpression and purification of the wild-type or mutant BmAEP1 zymogen

The DNA fragment encoding the transcriptomic sequencing-derived BmAEP1 zymogen (residues 22[484) was chemically synthesized at Tsingke Biotechnology (Beijing, China), and then ligated into a pET vector after cleavage with restriction enzymes NdeI and EcoRI. The resultant expression construct pET/6×His-BmAEP1 encodes an N-terminally 6×His-tagged BmAEP1 zymogen whose nucleotide and amino acid sequences are shown in Fig. S3. Site-directed mutagenesis of the wild-type BmAEP1 zymogen was conducted via the QuikChange approach using pET/6×His-BmAEP1 as a template.

For bacterial overexpression, the expression construct encoding the wild-type or mutant BmAEP1 zymogen was transformed into the *E. coli* strain Shuffle-T7. The transformed bacteria were cultured in liquid Luria-Bertani (LB) medium (plus 100 μg/ml ampicillin) to OD_600_≍1.0 at 37 °C with vigorous shaking (250 rpm). Thereafter, isopropyl β-D-thiogalactoside (IPTG) was added to a final concentration of 0.7 mM and the bacteria were continually cultured at 20 °C for ∼20 h with gentle shaking (100 rpm). After harvested by centrifugation (5000 *g*, 10 min), the bacteria were resuspended in lysis buffer (40 mM sodium phosphate, 100 mM NaCl, 0.1% Triton X-100, pH 7.0) and lysed by sonication. After centrifugation (12000 *g*, 30 min), the supernatant was applied to a Ni^2+^ column (Beyotime Biotechnology), and the recombinant zymogen was eluted by 100 mM imidazole. For the wild-type zymogen, the eluted fraction was dialyzed against ice-cold storage solution (20 mM phosphate, 100 mM NaCl, 20% sucrose, 1.0 mM EDTA, 5.0 mM β-mercaptoethanol, pH 6.5) for ∼2 h and stored at −80 °C. For the mutant zymogen, the eluted fraction was dialyzed against ice-cold cleavage buffer (20 mM phosphate, pH 7.0) for ∼2 h and stored at −80 °C. Samples at different preparation steps were analyzed by SDS-PAGE.

### 2.8. Self-activation of the wild-type BmAEP1 zymogen

To start self-activation, the recombinant wild-type BmAEP1 zymogen (∼3.5 μM) was mixed with equal volume of the activation buffer (200 mM sodium acetate, pH 4.4). After incubation at 25 °C for the indicated time, 10 μl of the reaction mixture was removed and diluted 40-fold using the neutralizing buffer (400 mM phosphate, pH 7.0) containing 0.2% BSA for subsequent peptide ligase activity assay. Alternatively, 14 μl of the reaction mixture was removed and mixed with 6 μl of SDS-gel loading buffer for subsequent SDS-PAGE analysis. For other activity assays, the reaction mixture was mixed with 2 volumes of the neutralizing buffer after self-activation for ∼60 min, put on ice, and used within a day.

### 2.9. Trypsin-assisted activation of [G238V]BmAEP1 zymogen

To activate [G238V]BmAEP1 zymogen, stock solution of bovine trypsin (Sigma-Aldrich, St. Louis, MO, USA) was added to the mutant zymogen fraction (∼1.0 μM) to a final concentration of 40 nM. After incubation at 20 °C for the indicated time, 14 μl of the reaction mixture was removed, mixed with 6 μl of the SDS-gel loading buffer, and analyzed by SDS-PAGE. After cleavage for 30 min, the reaction mixture was loaded onto a DEAE ion-exchange column. After thoroughly washing by 200 mM NaCl, the bound [G238V]BmAEP1 was eluted from the column by 400 mM NaCl (in 20 mM phosphate, pH 7.0). The eluted fraction was stored at −80 [after glycerol and sucrose were added to the final concentration of 20% and 15%, respectively.

To dissociate the noncovalently bound cap domain, the trypsin-treated [G238V]BmAEP1 (∼4 μM) was mixed with 1/5 volume of the activation buffer (200 mM acetate, pH 4.4). After incubation at 20 [for the indicated time, 10 μl of the reaction mixture was removed and diluted 330-fold using the neutralizing buffer (200 mM phosphate, pH 7.0) containing 0.2% BSA for subsequent peptide ligase activity assay. To prepare the acidic activated enzyme for other assays, the activation mixture was mixed with 2/3 volume of the neutralizing buffer without BSA, put on ice, and used within a day.

### 2.10. Ligation of the NanoLuc reporter with peptides by [G238V]BmAEP1

The purified ligation version NanoLuc reporter (NanoLuc-NHV or NanoLuc-NAL), the synthetic ligation version peptide (GI-SmBiT or GI-SP), and the activated [G238V]BmAEP1 were mixed together in 200 mM phosphate buffer (pH 6.5) at the indicated final concentrations. After incubation at 20 °C for the indicated time, 10 μl of the reaction mixture was removed and mixed with 40 μl of SDS-gel loading buffer. After boiling, 10 μl of the boiled sample was loaded onto a 15% SDS-gel for electrophoresis. To prepare the NanoLuc-ligated SP tracer (Luc-SP) for receptor binding assay, the reaction mixture was subjected to a spin Ni^2+^ column after ligation for 2 h. After washing off the excess amount of GI-SP, the expected ligation product was eluted by 250 mM imidazole, and then analyzed by SDS-PAGE and bioluminescence measurement.

### 2.11. Cyclization of the synthetic SFTI-NHV by [G238V]BmAEP1

The lyophilized synthetic SFTI-NHV was weighed using a precise balance and dissolved in 1.0 mM aqueous HCl (pH 3.0) as a stock solution (25 mM). For oxidative refolding, the peptide stock solution was diluted into cold refolding buffer (0.5 M L-arginine, pH 9.0, plus 90 μM 2,2’-dithiodipyridine) to a final concentration of 30 μM. After incubation on ice for 1.5 h, the refolding mixture was acidified to pH 3[4 and applied to HPLC. The refolded SFTI-NHV was eluted from the C_18_ reverse-phase column (Zorbax 300SB-C18, 9.4 or 4.6 × 250 mm; Agilent Technologies) by an acidic acetonitrile gradient, manually collected, and lyophilized.

For cyclization, the refolded SFTI-NHV and the activated [G238V]BmAEP1 were added to the ligation buffer (200 mM phosphate, pH 7.0) to the final concentration of 120 μM and 120 nM, respectively. After ligation at 20 [for the indicated time, 100 μl of the reaction mixture was removed, acidified to pH 3[4, and analyzed by HPLC using a C_18_ reverse-column (Zorbax 300SB-C18, 4.6 × 250 mm; Agilent Technologies). Molecular mass of the refolded SFTI-NHV before or after cyclization was measured by electrospray mass spectrometry.

### 2.12. Receptor binding assay of the Luc-SP tracer

Human embryonic kidney (HEK) 293T cells were transfected with the expression construct pENTER/TACR1 (WZ Bio, Jinan, Shandong Province, China) using the transfection reagent Lipo 8000 (Beyotime Biotechnology). Next day, the transfected cells were trypsinized, seeded into a 96-well white plate, and continuously cultured for ∼24 h to ∼100% confluence. To conduct the saturation binding assay, the medium was removed, and the binding solution (DMEM plus 1% BSA) containing varied concentrations of Luc-SP with or without 1.0 μM of synthetic GI-SP was added (50 μl/well). After incubation at 22 °C for 1 h, the binding solution was removed and the cells were washed twice with ice-cold PBS (200 μl/well/time). Thereafter, diluted NanoLuc substrate was added (50 μl/well) and bioluminescence was immediately measured on a SpectraMax iD3 plate reader (Molecular Devices).

The measured bioluminescence data were expressed as mean ± SD (*n* = 3) and fitted to one-site binding model using SigmaPlot10.0 software. Amount of the receptor-bound tracer was calculated from the measured specific binding according to the measured specific luciferase activity of the Luc-SP tracer.

## 3. Results

### 3.1. AEP-type peptide ligase activity is widely present in bamboo species

In the present study, we screened 61 bamboo species from 16 genera using the newly developed NanoBiT-based peptide ligase activity assay (Table 1). After the crude extract of fresh bamboo leaves was mixed with peptide ligase substrates, recombinant LgBiT-NHV and synthetic GI-SmBiT, increased bioluminescence was detected in all samples, although the measured bioluminescence varied significantly among different bamboo species (Table 1). In longer time assays, the measured bioluminescence increased quickly after mixing the crude extract with both substrates; however no increase in bioluminescence was detected in the absence of GI-SmBiT (Fig. S4). Thus, it seemed that AEP-type peptide ligase activity is widely present in bamboo species. To the best of our knowledge, this was the first time that AEP-type peptide ligase activity was detected in so many phylogenetically related higher plants.

**Table 1.** AEP-type peptide ligase activity detected in the crude extract of bamboo leaves. To start the peptide ligase activity assay, the crude extract was mixed with equal volume of substrates mixture (10 μM LgBiT-NHV and 20 μM GI-SmBiT). After incubation at 20 °C for 10 min, 10 μl of the ligation mixture was removed and diluted 2,000-fold using NanoBiT diluting solution. Finally, the diluted mixture was transferred to a 384-well white plate (10 μl/well) and bioluminescence was immediately measured after addition of the diluted NanoLuc substrate (10 μl/well). The measured bioluminescence data are expressed as mean ± SD (*n* = 2[3). Concentrations of the accumulated ligation product were calculated from the measured bioluminescence and the measured specific activity of the HiBiT-complemented LgBiT-NHV.

| Number | Bamboo species | Bioluminescence (RLU) | Product (μM) |
| --- | --- | --- | --- |
| 1 | *Bambusa multiplex* 'Alphonse-Karri' | 405585 ± 18104 | 0.27 ± 0.01 |
| 2 | *Bambusa multiplex* f. fernleaf | 361371 ± 18357 | 0.24 ± 0.01 |
| 3 | *Bambusa multiplex* | 371781 ± 28076 | 0.25 ± 0.02 |
| 4 | *Bambusa ventricosa* | 392415 ± 8118 | 0.26 ± 0.01 |
| 5 | *Bambusa ventricosa* cv. Kimmei | 337530 ± 23907 | 0.23 ± 0.02 |
| 6 | *Brachystachyum densiflorum* | 406724 ± 29863 | 0.27 ± 0.02 |
| 7 | *Chimonobambusa neopurpurea* | 224139 ± 14283 | 0.15 ± 0.01 |
| 8 | *Drepanostachyum scandens* | 659653 ± 55946 | 0.44 ± 0.04 |
| 9 | *Hibanobambus tranquillans* | 114947 ± 1380 | 0.08 ± 0.00 |
| 10 | *Indocalamus herklotsii* | 69497 ± 6278 | 0.05 ± 0.00 |
| 11 | *Indocalamus latifolius* | 246068 ± 18121 | 0.16 ± 0.01 |
| 12 | *Indocalamus tessellatus* | 281906 ± 34337 | 0.19 ± 0.02 |
| 13 | *Indosasa shibataeoides* | 509115 ± 6674 | 0.44 ± 0.00 |
| 14 | *Indosasa sinica* | 99571 ± 7462 | 0.07 ± 0.00 |
| 15 | *Oligostachyum lubricum* | 316959 ± 26139 | 0.21 ± 0.02 |
| 16 | *Oligostachyum sulcatum* | 263918 ± 1958 | 0.18 ± 0.00 |
| 17 | *Phyllostachys aurea* | 340120 ± 25279 | 0.23 ± 0.02 |
| 18 | *Phyllostachys aureosulcata* cv. Aureocarlis | 371683 ± 161 | 0.25 ± 0.00 |
| 19 | *Phyllostachys aureosuleata* cv. Pekinensis | 354240 ± 13147 | 0.24 ± 0.01 |
| 20 | *Phyllostachys aureosulcata* 'Spectabilis' | 512932 ± 45799 | 0.34 ± 0.03 |
| 21 | *Phyllostachys bambusoides* f. lacrima-deae | 489503 ± 49580 | 0.33 ± 0.03 |
| 22 | *Phyllostachys bambusoides* f. duihuazhu | 411126 ± 5065 | 0.27 ± 0.00 |
| 23 | *Phyllostachys edulis* | 194759 ± 9091 | 0.13 ± 0.01 |
| 24 | *Phyllostachys edulis* cv. Pachyloen | 314475 ± 31009 | 0.21 ± 0.02 |
| 25 | *Phyllostachys flexuosa* | 294460 ± 4462 | 0.20 ± 0.00 |
| 26 | *Phyllostachys glauca* | 150144 ± 8477 | 0.10 ± 0.01 |
| 27 | *Phyllostachys glauca* var. *variabilis* | 496853 ± 42311 | 0.33 ± 0.03 |
| 28 | *Phyllostachys heteroclada* | 525165 ± 20697 | 0.35 ± 0.01 |
| 29 | *Phyllostachys heterocycla* | 192529 ± 12200 | 0.13 ± 0.01 |
| 30 | *Phyllostachys heterocycla* cv. Tao-Kiang | 47476 ± 5211 | 0.03 ± 0.00 |
| 31 | *Phyllostachys heterocycle* f. *gracilis* | 410596 ± 10226 | 0.27 ± 0.01 |
| 32 | *Phyllostachys heterocycla* cv. Luteosulcata | 502145 ± 28960 | 0.33 ± 0.02 |
| 33 | *Phyllostachys incarnata* | 304642 ± 4919 | 0.20 ± 0.00 |
| 34 | *Phyllostachys iridescens* | 367839 ± 42586 | 0.25 ± 0.03 |
| 35 | *Phyllostachys kwangsiensis* | 299069 ± 19702 | 0.20 ± 0.01 |
| 36 | *Phyllostachys nidularia* f. *mirabilis* | 384270 ± 46507 | 0.26 ± 0.03 |
| 37 | *Phyllostachys nidularia* | 393944 ± 37029 | 0.26 ± 0.02 |
| 38 | *Phyllostachys nigra* | 470176 ± 60703 | 0.31 ± 0.04 |
| 39 | *Phyllostachys nigra* var. *henonis* | 355888 ± 23229 | 0.24 ± 0.02 |
| 40 | *Phyllostachys nuda* f. *lacalis* | 195506 ± 2321 | 0.13 ± 0.00 |
| 41 | *Phyllostachys nuda* | 306113 ± 14404 | 0.20 ± 0.01 |
| 42 | *Phyllostachys praecox* cv. 'Praecox' | 252318 ± 14342 | 0.17 ± 0.01 |
| 43 | *Phyllostachys prominens* | 162530 ± 2505 | 0.11 ± 0.00 |
| 44 | *Phyllostachys edulis* f. *tubaeformis* | 313385 ± 11638 | 0.21 ± 0.01 |
| 45 | *Pleioblastus amarus* | 154210 ± 9422 | 0.10 ± 0.01 |
| 46 | *Pleioblastus china* f. *hisauchii* | 252558 ± 3290 | 0.17 ± 0.00 |
| 47 | *Pleioblastus gramineus* | 147360 ± 1407 | 0.10 ± 0.00 |
| 48 | *Pleioblastus kongosanensis* | 223912 ± 10733 | 0.15 ± 0.01 |
| 49 | *Pleioblastus simonii* f. *heterophyllus* | 127053 ± 9096 | 0.08 ± 0.01 |
| 50 | *Pseudosasa amabilis* | 292179 ± 13468 | 0.19 ± 0.01 |
| 51 | *Pseudosasa japonica* | 125436 ± 16231 | 0.08 ± 0.01 |
| 52 | *Pseudosasa japonica* var. *tsutsumiana* | 182747 ± 13652 | 0.12 ± 0.01 |
| 53 | *Qiongzhusa tumidinoda* | 230159 ± 10393 | 0.15 ± 0.01 |
| 54 | *Sasa argenteistriatus* | 178438 ± 10384 | 0.12 ± 0.01 |
| 55 | *Sasa fortunei* | 263789 ± 6713 | 0.18 ± 0.00 |
| 56 | *Sasa viridistriatus* f. *chrysophyllus* | 142365 ± 794 | 0.10 ± 0.00 |
| 57 | *Semiarundinaria fastuosa* | 368813 ± 33708 | 0.25 ± 0.02 |
| 58 | *Semiarundinaria sinica* | 113373 ± 14303 | 0.08 ± 0.01 |
| 59 | *Shibataea chinensis* | 304308 ± 22430 | 0.20 ± 0.02 |
| 60 | *Sinobambusa parvifolia* | 358545 ± 30919 | 0.24 ± 0.02 |
| 61 | *Sinobambusa tootsik* | 147441 ± 9009 | 0.10 ± 0.01 |

### 3.2. Identification of an AEP-type peptide ligase candidate from Bambusa multiplex

To identify the peptide ligase from bamboo, fresh leaves of *Bambusa multiplex* were subjected to transcriptomic sequencing, because this species displayed high peptide ligase activity (Table 1 and Fig. S4) and is a popular horticultural bamboo species in China. From the assembled transcriptomic sequencing data, a full-length pre-zymogen of a possible AEP-type peptide ligase (BmAEP1) was identified according to its sequence homology with butelase-1 (Fig. 1A and Fig. S2). The full-length BmAEP1 pre-zymogen shares a common architecture and considerable sequence homology with the pre-zymogens of some AEP-type peptide ligases, including an N-terminal signal peptide, a short N-terminal domain, a catalytic core domain, a short linker, and a C-terminal cap domain (Fig. 1A).

**Fig. 1.**
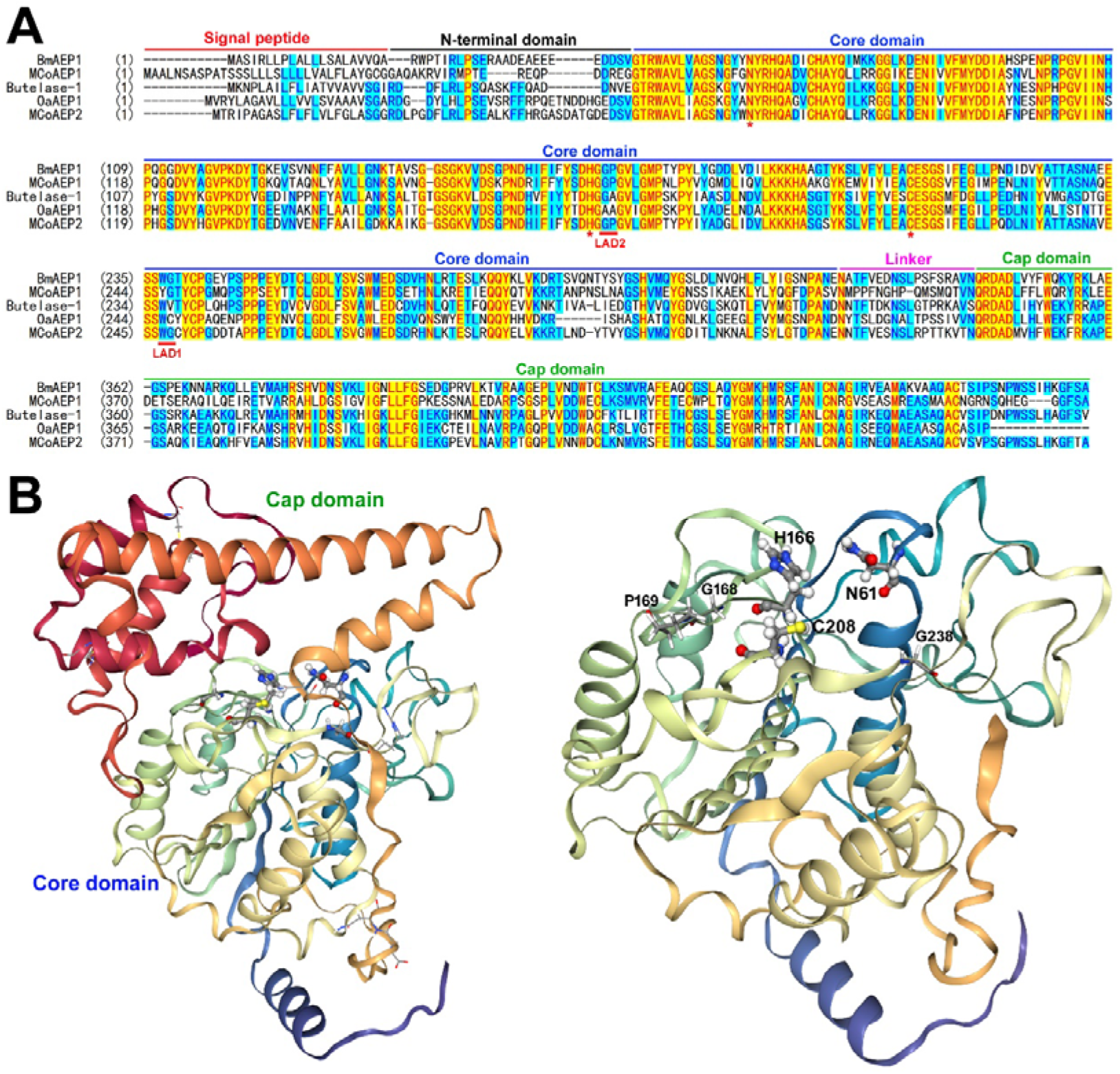
BmAEP1 identified from *Bambusa multiplex.* (**A**) Amino acid sequence alignment of the transcriptomic sequencing-derived BmAEP1 pre-zymogen with pre-zymogens of some AEP-type peptide ligases. The residues forming the catalytic triad are indicated by asterisks. (**B**) Structure of BmAEP1 zymogen predicted by RoseTTAFold algorithm. Left panel, the overall structure of the intact zymogen; right panel, the catalytic core domain. The residues forming the catalytic triad are shown as sticks-and-balls, and those in ligase activity determinants (LAD1 and LAD2) are shown as sticks.

The presence of the BmAEP1 pre-zymogen was confirmed by cDNA cloning. After reverse transcription, PCR amplification, and cloning, two full-length BmAEP1 cDNAs were identified from three transformed *E. coli* colonies: one cDNA had a missense mutation (Y297L) and two synonymous mutations, the other had two missense mutations (H95Y and Y297L) and nine synonymous mutations compared with the transcriptomic sequencing-derived BmAEP1 pre-zymogen (Fig. S2).

The RoseTTAFold algorithm predicted that the BmAEP1 zymogen (residues 22[484) shares a similar structure with other AEP zymogens (Fig. 1B). Its C-terminal cap domain covers the substrate binding site of the catalytic core domain (Fig. 1B, left panel); therefore, the zymogen was expected to be inactive. The cap domain was stabilized by two disulfide bonds, Cys432[Cys467 and Cys420[Cys450. After removal of the cap domain by cleavage at the linker region, the catalytic center composed of Cys208, His166, and Asn61 would be exposed and the free core domain would restore the catalytic activity (Fig. 1B, right panel).

### 3.3. Bacterial overexpression, self-activation, and enzymatic properties of the *wild-type BmAEP1*

To test whether BmAEP1 is a peptide ligase, an N-terminally 6×His-tagged BmAEP1 zymogen (residues 22[484) was overexpressed in the *E. coli* strain Shuffle-T7, because this strain can promote disulfide formation in the cytosol. After purification using a Ni^2+^ column, a major band slightly lower than 60 kDa was eluted from the column, as analyzed by SDS-PAGE (Fig. 2A). This band was expected to be the intact 6×His-BmAEP1 zymogen whose theoretical molecular weight was 52.3 kDa. After purification, 2−3 mg of the recombinant zymogen was typically obtained from 1 liter of *E. coli* culture broth.

**Fig. 2.**
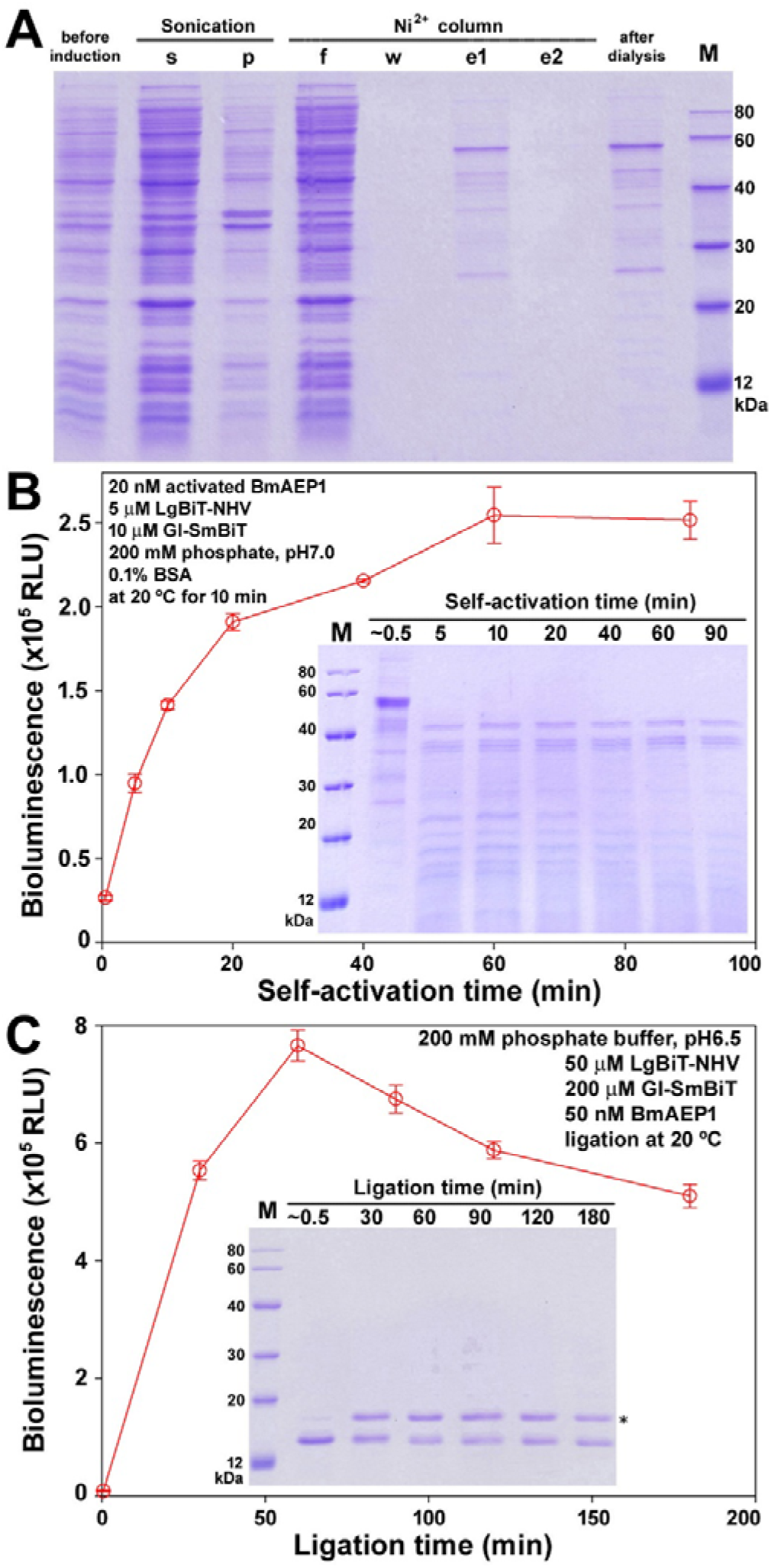
Bacterial overexpression, self-activation, and ligation property of the wild-type BmAEP1. (**A**) SDS-PAGE analysis of the recombinant wild-type BmAEP1 zymogen at different preparation steps. Samples at the indicated steps were loaded onto a 15% SDS-gel and the gel was stained by Coomassie brilliant blue 250 after electrophoresis. Lane (before induction), bacteria lysate before IPTG induction; lane (s), supernatant; lane (p), pellet; lane (f), flow-through; lane (w), washing fraction by 20 mM imidazole; lane (e1), eluted fraction by 100 mM imidazole; lane (e2), eluted fraction by 500 mM imidazole; lane (after dialysis), after dialysis of the fraction eluted by 100 mM imidazole. (**B**) Self-activation of the wild-type BmAEP1 zymogen monitored by peptide ligase activity assay and SDS-PAGE. To start self-activation, the purified BmAEP1 zymogen (∼3.5 μM) was mixed with equal volume of the activation buffer (200 mM sodium acetate, pH 4.4). After incubation at 25 °C for the indicated time, 10 μl of the reaction mixture was removed and diluted 40-fold using the neutralizing buffer (200 mM phosphate, pH 7.0) with 0.2% BSA. Thereafter, 10 μl of the diluted enzyme was mixed with equal volume of the substrate mixture (10 μM LgBiT-NHV and 20 μM GI-SmBiT) to start the ligase activity assay. After incubation at 20 °C for 10 min, 10 μl of the ligation mixture was removed and diluted 2,000-fold using NanoBiT diluting solution for subsequent bioluminescence measurement. **Inner panel**, SDS-PAGE analysis. After the zymogen was self-activated at 25 °C for the indicated time, 14 μl of the reaction mixture was removed and mixed with 6 μl of SDS-gel loading buffer. After boiling, the samples were loaded onto a 15% SDS-gel and the gel was stained by Coomassie brilliant blue R250 after electrophoresis. (**C**) Wild-type BmAEP1-catalyzed ligation of LgBiT-NHV and GI-SmBiT monitored by bioluminescence and SDS-PAGE. To start ligation, self-activated BmAEP1 was mixed with substrates at the indicated concentrations. After ligation at 20 °C for the indicated time, 10 μl of the reaction mixture was removed and diluted 100,000-fold using NanoBiT diluting solution for bioluminescence measurement. **Inner panel**, SDS-PAGE analysis. At the indicated time, 10 μl of the ligation mixture was removed and mixed with 40 μl of SDS-gel loading buffer. After boiling, 10 μl of the boiled mixture was loaded onto a 15% SDS-gel and the gel was stained by Coomassie brilliant blue 250 after electrophoresis.

After elution from the Ni^2+^ column, the zymogen fraction was subjected to dialysis to remove imidazole because it inhibits self-activation of the zymogen. After dialysis, the zymogen could be stored at −80 °C for several months without significant inactivation. For convenient self-activation, we developed a simple procedure via mixing the zymogen stock with acidic activation buffer. SDS-PAGE analysis showed that the ∼60 kDa zymogen band was quickly converted to several smaller bands around 40 kDa after self-activation (Fig. 2B, inner panel), suggesting that the BmAEP1 zymogen can cleave itself efficiently under acidic conditions. In the peptide ligase activity assay using LgBiT-NHV and GI-SmBiT as substrates, the measured bioluminescence increased slowly after self-activation (Fig. 2B), suggesting that dissociation of the non-covalently bound cap domain from the core domain is a slow process after self-cleavage of the zymogen.

Subsequently, we analyzed the ligation property of the wild-type BmAEP1 using LgBiT-NHV and GI-SmBiT as substrates. After 50 μM of LgBiT-NHV was ligated with 200 μM GI-SmBiT by 50 nM of BmAEP1, the measured bioluminescence increased quickly within 60 min, but then decreased slowly (Fig. 2C). As calculated from the measured bioluminescence and the measured specific activity of the HiBiT-complemented LgBiT-NHV, at most 60% of LgBiT-NHV was converted to the ligation product. SDS-PAGE analysis showed that a slightly larger band (indicated by an asterisk), presumably the ligation product, appeared quickly after the initiation of ligation (Fig. 2C, inner panel); however, its band density decreased slowly after long term incubation, implying that BmAEP1 can hydrolyze the ligation product via its intrinsic hydrolysis activity. Thus, it seemed that the wild-type BmAEP1 has high peptide ligase activity, but with considerable hydrolysis activity.

### 3.4. Bacterial overexpression and trypsin-assisted activation of [G238V]BmAEP1 zymogen

To eliminate the intrinsic hydrolysis activity of the wild-type BmAEP1, we conducted site-directed mutagenesis at its ligase activity determinants, LAD1 and LAD2 (Fig. 1A). Mutations at LAD2 (Gly168Ala or Pro169Ala) drastically decreased its expression level in *E. coli*, whereas mutation at LAD1 (Gly238Val) had no significant effect on its expression in *E. coli*. Thus, we prepared the [G238V]BmAEP1 zymogen for enzymatic property studies.

After the [G238V]BmAEP1 zymogen was overexpressed in Shuffle-T7 and purified via a Ni^2+^ column, a major band slightly lower than 60 kDa appeared on SDS-PAGE (Fig. 3A). However, the mutant zymogen failed to self-cleave after dialysis, despite trying various conditions. For self-activation, the AEP zymogens cleave peptide bonds at the C-terminus of Asn or Asp residues in their linker region (Fig. 1A). We found that there is an Arg residue in the linker region of the BmAEP1 zymogen (Fig. 1A), and thus speculated whether trypsin could be used to activate the mutant zymogen. Trypsin is a commercially available and cheap endoproteinase that specifically cleaves the peptide bond at the C-terminus of a positively charged Arg or Lys residue. Moreover, trypsin is a basic protein with an isoelectric point of ∼10, thus it can be easily separated from the acidic AEPs after cleavage using an ion-exchange chromatography.

**Fig. 3.**
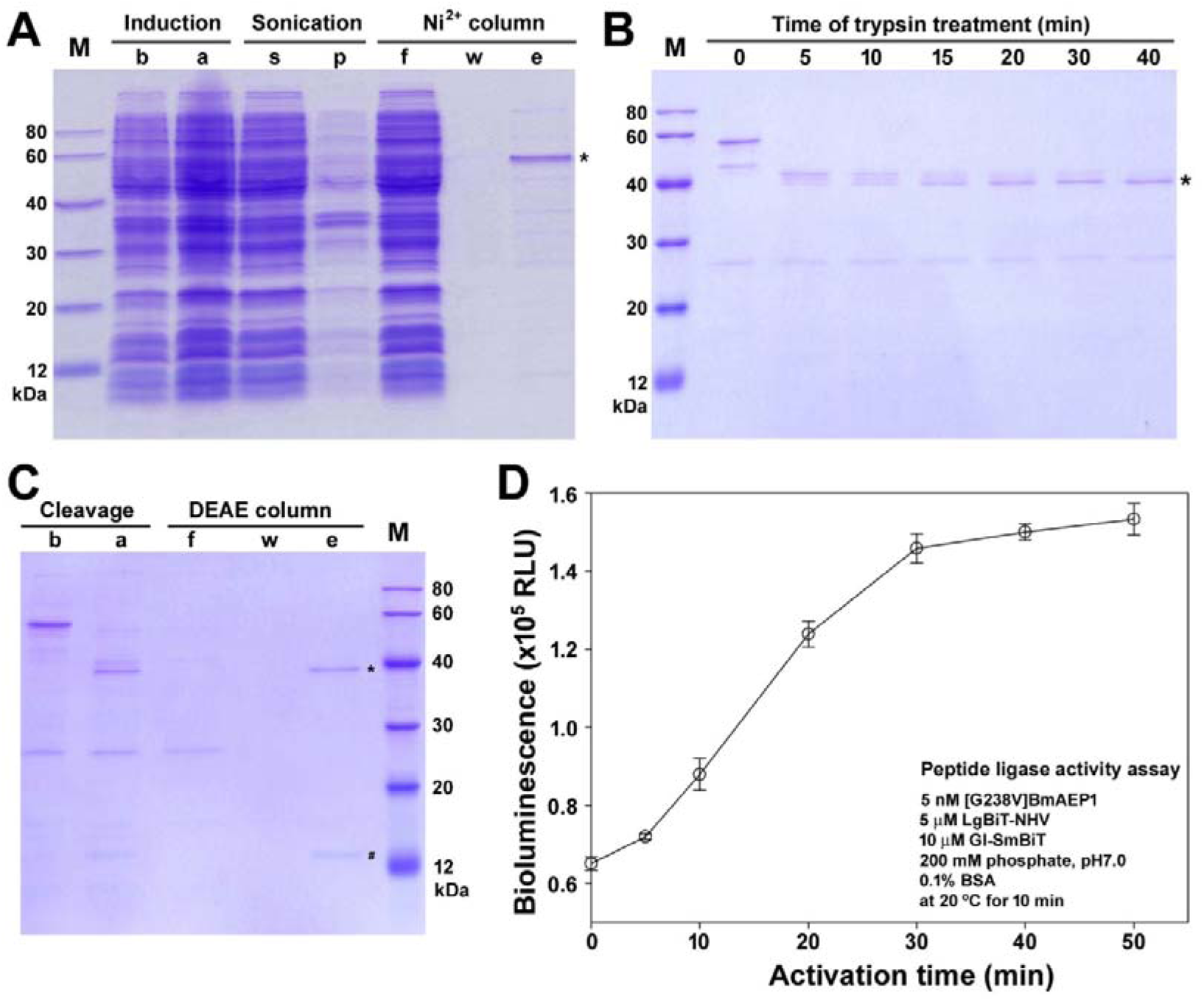
Bacterial overexpression and trypsin-assisted activation of [G238V]BmAEP1 zymogen. (**A**) SDS-PAGE analysis of the recombinant [G238V]BmAEP1 zymogen at different preparation steps. Samples at the indicated stages were loaded onto a 15% SDS-gel and the gel was stained by Coomassie brilliant blue 250 after electrophoresis. Lane (b), before IPTG induction; lane (a), after IPTG induction; lane (s) supernatant; lane (p), pellet; lane (f), flow-through; lane (w), washing fraction by 20 mM imidazole; lane (e), eluted fraction by 100 mM imidazole. (**B**) SDS-PAGE analysis of the trypsin-assisted activation of [G238V]BmAEP1 zymogen. After the purified zymogen (∼1.0 μM) was treated with bovine trypsin (40 nM) at 20 °C for the indicated time, 14 μl of the reaction mixture was removed, mixed with 6 μl of SDS-gel loading buffer, and loaded onto a 15% SDS-gel after boiling. After electrophoresis, the gel was stained by Coomassie brilliant blue 250. (**C**) SDS-PAGE analysis of the trypsin cleaved [G238V]BmAEP1 zymogen after ion-exchange chromatography. Lane (b), before trypsin treatment; lane (a), after trypsin treatment for 30 min; lane (f), flow-through fraction; lane (w), washing fraction by 200 mM NaCl; lane (e), eluted traction by 400 mM NaCl. Samples at the indicated stages were loaded onto a 15% SDS-gel and the gel was stained by Coomassie brilliant blue 250 after electrophoresis. (**D**) Peptide ligase activity assay of [G238V]BmAEP1 after acidic activation. For acidic activation, the trypsin-treated [G238V]BmAEP1 fraction (∼4 μM) eluted from the DEAE column was mixed with 1/5 volume of the activation buffer (200 mM sodium acetate, pH 4.4). After incubation at 20 °C for the indicated time, 10 μl of the mixture was removed and diluted 330-fold using the neutralizing buffer (200 mM phosphate, pH 7.0) with 0.2% BSA. Thereafter, the diluted enzyme was mixed with equal volume of substrate mixture (10 μM LgBiT-NHV and 20 μM GI-SmBiT) to start the peptide ligase activity assay. After incubation at 20 °C for 10 min, 10 μl of the ligation mixture was removed and diluted 2,000-fold using NanoBiT diluting solution for subsequent bioluminescence measurement.

After optimization, bovine trypsin could efficiently cleave the [G238V]BmAEP1 zymogen, as analyzed by SDS-PAGE (Fig. 3B): the ∼60 kDa zymogen band was quickly converted to three smaller bands of ∼40 kDa. After cleavage, the reaction mixture was subjected to an anion ion-exchange chromatography: the basic trypsin flowed through the DEAE column, and then the bound [G238V]BmAEP1 was eluted. SDS-PAGE analysis (Fig. 3C) showed that the eluted fraction contained an ∼40 kDa band (indicated by an asterisk), presumably the catalytic core domain, as well as an ∼12 kDa band (indicated by an octothorpe), presumably the cleaved cap domain, suggesting the cap domain was likely to be non-covalently bound to the core domain after trypsin cleavage. The trypsin-treated [G238V]BmAEP1 fraction could be store at −80 °C for several months without significant inactivation.

To dissociate the non-covalently bound cap domain, the eluted fraction was subjected to acidic activation, which indeed increased the peptide ligase activity, as shown in Fig. 3D. Moreover, the eluted fraction had considerable peptide ligase activity before acidic activation (Fig. 3D), implying that non-covalently bound cap domain could slowly dissociate from the core domain even without acidic activation.

### 3.5. Ligation of LgBiT-NHV with GI-SmBiT by [G238V]BmAEP1

We first tested the effect of pH on the ligation rate and product accumulation of [G238V]BmAEP1 using LgBiT-NHV and GI-SmBiT as substrates (Fig. 4A[C). According to the measured bioluminescence, [G238V]BmAEP1 displayed the highest initial ligation rate at pH 6.5 (Fig. 4A), and also accumulated the most ligation product at this pH value (Fig. 4B). Based on the measured bioluminescence and measured specific activity of the HiBiT-complemented LgBiT-NHV, ∼95% of LgBiT-NHV was converted to the ligation product at pH 6.5 after 3 h. The presence of the expected ligation product was confirmed by SDS-PAGE (Fig. 4C): the band of LgBiT-NHV was converted to a slightly larger band, presumably the ligation product. As estimated from the band density, over 90% of LgBiT-NHV was converted to the ligation product at pH 6.0[7.0. Thus, it seemed that [G238V]BmAEP1 works well at pH 6.0[7.0 for ligation of LgBiT-NHV and GI-SmBiT.

**Fig. 4.**
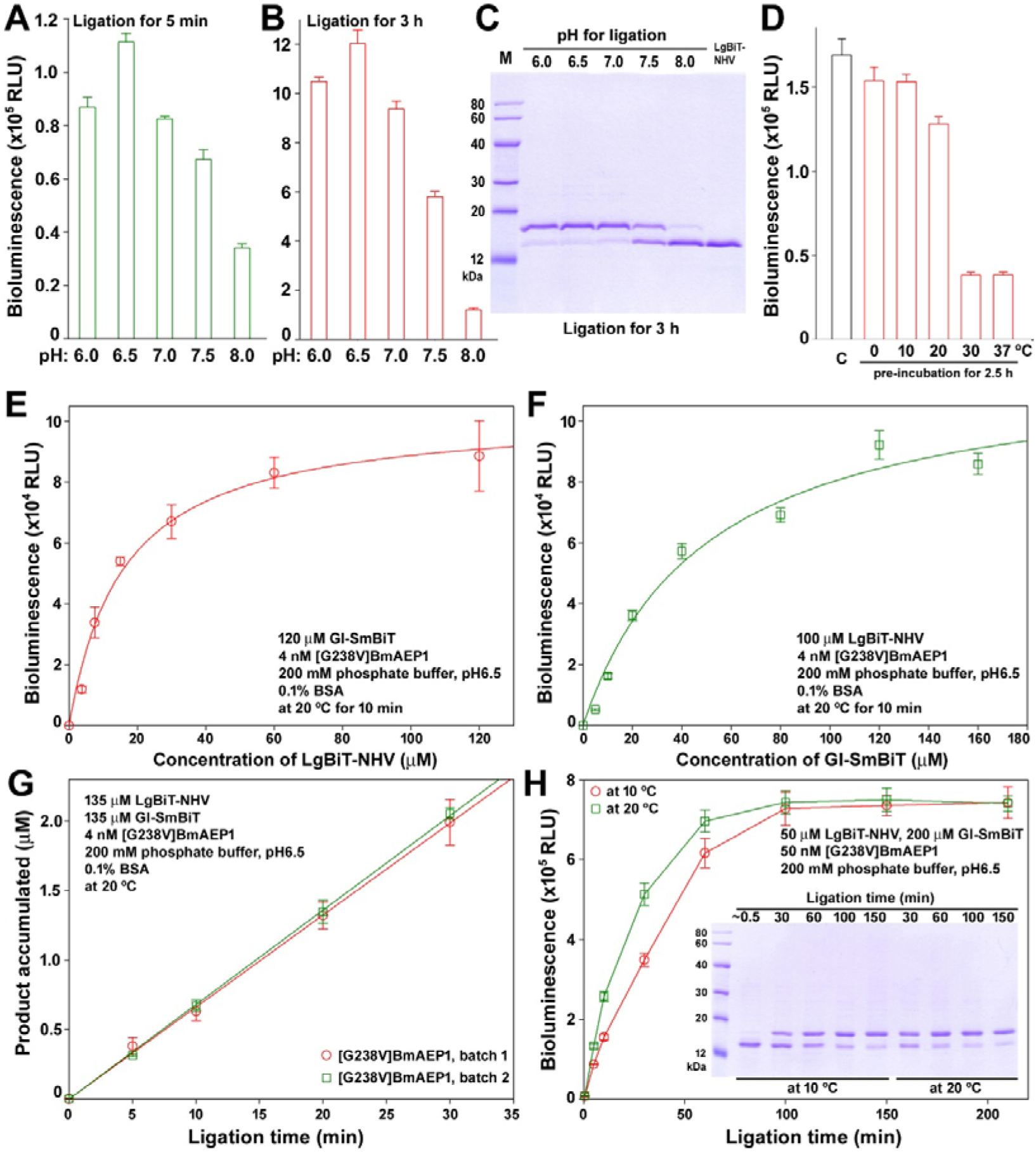
Ligation of LgBiT-NHV and GI-SmBiT by [G238V]BmAEP1. (**A**□**C**) Effect of pH on initial ligation velocity (**A**) and ligation product accumulation (**B,C**) analyzed by bioluminescence (**A,B**) and SDS-PAGE (**C**). 50 μM of LgBiT-NHV was ligated with 200 μM of GI-SmBiT by 50 nM of [G238V]BmAEP1 in 200 mM phosphate buffer with different pH values. After ligation at 20 °C for 5 min or for 3 h, 10 μl of the reaction mixture was removed and diluted 100,000-fold using NanoBiT diluting solution for bioluminescence measurement. For SDS-PAGE analysis, 10 μl of the reaction mixture was removed after ligation at 20 °C for 3 h, and mixed with 40 μl of SDS-gel loading buffer. After boiling, 10 μl of the mixture was loaded onto a 15% SDS-gel and the gel was stained by Coomassie brilliant blue 250 after electrophoresis. (**D**) Thermal stability of [G238V]BmAEP1. The enzyme was diluted into the ligation buffer (200 mM phosphate, pH 6.5, plus 0.1% BSA) to 10 nM, and then incubated at the indicated temperature for 2.5 h or not (control “C”). Thereafter, [G238V]BmAEP1 was mixed with equal volume of substrate mixture (10 μM LgBiT-NHV and 20 μM GI-SmBiT). After incubation at 20 °C for 10 min, the ligation mixture was diluted 2,000-fold using NanoBiT diluting solution for bioluminescence measurement. (**E,F**) Enzymatic kinetics of [G238V]BmAEP1 towards substrate LgBiT-NHV (**E**) and GI-SmBiT (**F**). Substrates and enzyme were mixed together in 200 mM phosphate buffer (pH 6.5) with 0.1% BSA at the indicated concentrations. After incubation at 20°C for 10 min, 10 μl of the reaction mixture was removed and diluted 10,000-fold using NanoBiT diluting solution for bioluminescence measurement. The measured bioluminescence data were fitted to the function of V = V_max_[S]/(K_m_+[S]) using SigmaPlot10.0 software. (**G**) Measurement of k_cat_ using two batches of [G238V]BmAEP1. Substrates and enzyme were mixed together in 200 mM phosphate buffer (pH 6.5) with 0.1% BSA at the indicated concentrations. After incubation at 20 °C for the indicated time, 10 μl of the ligation mixture was removed and diluted 100,000-fold using NanoBiT diluting solution for bioluminescence measurement. (**H**) Time course of the [G238V]BmAEP1-catalyzed ligation of LgBiT-NHV and GI-SmBiT monitored by bioluminescence and SDS-PAGE. Substrates and enzyme were mixed together in 200 mM phosphate buffer (pH 6.5) at the indicated concentrations. After incubation at 20 °C or 10 °C for the indicated time, 10 μl of the ligation mixture was removed and diluted 100,000-fold using NanoBiT diluting solution for bioluminescence measurement. **Inner panel**, SDS-PAGE analysis. At the indicated times, 10 μl of the ligation mixture was removed and mixed with 40 μl of SDS-gel loading buffer. After boiling, 10 μl of the boiled samples was loaded onto a 15% SDS-gel and the gel was stained by Coomassie brilliant blue 250 after electrophoresis.

After pre-incubation at 20 °C or a lower temperature for 2.5 h, [G238V]BmAEP1 retained over 75% of its activity, but pre-incubation at 30 or 37 °C for 2.5 h caused ∼80% activity loss (Fig. 4D). Thus, it seemed that [G238V]BmAEP1 is not so stable at high temperature. Based on its stability, we conducted ligation experiments at 20 °C or lower.

Thereafter, we determined the K_m_ values of [G238V]BmAEP1 towards LgBiT-NHV and GI-SmBiT. According to the measured bioluminescence, hyperbolic saturation curves were obtained for both substrates (Fig. 4E,F), with calculated K_m_ values of 15.9 ± 2.1 μM for LgBiT-NHV and 49.5 ± 6.5 μM for GI-SmBiT. At the saturated concentration of both substrates (135 μM), two batches of [G238V]BmAEP1 gave a consistent k_cat_ value of ∼17 min^-1^ for the ligation of LgBiT-HNV to GI-SmBiT (Fig. 4G). Thus, it seemed that [G238V]BmAEP1 could efficiently catalyze inter-molecular protein ligation.

Finally, we analyzed the ligation process via bioluminescence and SDS-PAGE (Fig. 4H). After 50 μM of LgBiT-NHV was ligated to 200 μM of GI-SmBiT by 50 nM of [G238V]BmAEP1, the measured bioluminescence increased quickly and reached a plateau at ∼1 h (Fig. 4H). The ligation rate at 10 °C was only slightly lower than that at 20 °C. According to the measured bioluminescence and the measured specific activity of HiBiT-complemented LgBiT-NHV, ∼95% of LgBiT-NHV could be finally converted to the ligation product. SDS-PAGE analysis confirmed the efficient ligation (Fig. 4H, inner panel). Moreover, long term incubation did not cause a decrease in bioluminescence and the band density of the ligation product (Fig. 4H), suggesting that the hydrolysis activity of [G238V]BmAEP1 is very low. Thus, the introduction of a mutation at LAD1 drastically reduced the intrinsic hydrolysis activity of BmAEP1.

### 3.6. Ligation of the NanoLuc reporter with peptides by [G238V]BmAEP1

The newly developed NanoLuc luciferase is the smallest and brightest luciferase reporter [41]. In recent years, our laboratory developed NanoLuc-based bioluminescent tracers for ligand[receptor binding assays by the covalent attachment of NanoLuc to an appropriate site of protein and peptide ligands via chemical or enzymatic approaches [42−47]. In the present study, we tested whether [G238V]BmAEP1 could be used to prepare NanoLuc-based bioluminescent tracers. For this purpose, we introduced an NHV or an NAL sequence to the C-terminus of a recombinant NanoLuc reporter (Fig. S1).

We first tested the ligation efficiency of [G238V]BmAEP1 towards the ligation version NanoLuc reporters (NanoLuc-NHV and NanoLuc-NAL) using GI-SmBiT as a nucleophilic peptide. As analyzed by SDS-PAGE, both proteins were quickly converted to a slightly larger band (presumably the ligation product, indicated by an asterisk in the figure) in pH 6.5 phosphate buffer (Fig. 5A). As estimated from the band density, the yield was ∼90% (Fig. 5A). Additionally, NanoLuc-NAL was slightly more suitable than NanoLuc-NHV as a substrate for [G238V]BmAEP1 (Fig. 5A). Besides the expected ligation product, there were a slightly smaller band (presumably a hydrolysis product, indicated by a letter “h” in the figure) and a much larger band (presumably the NanoLuc dimer, indicated by a letter “d” in the figure), but their percentage was quite low (Fig. 5A). Thus, it seemed that [G238V]BmAEP1 could efficiently catalyze the ligation of the NanoLuc reporter to peptides.

**Fig. 5.**
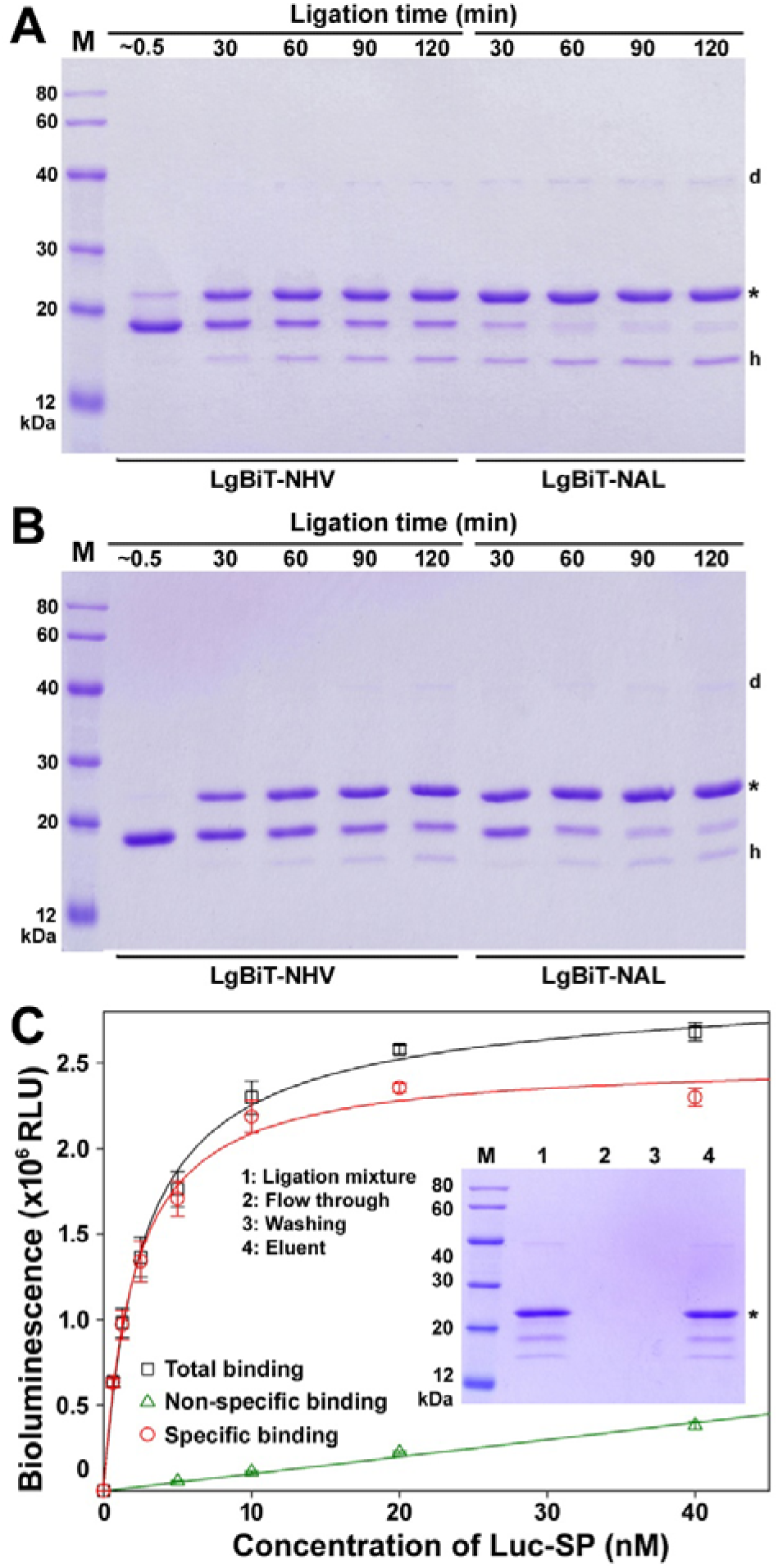
Ligation of the NanoLuc reporter with peptides by [G238V]BmAEP1. (**A,B**) SDS-PAGE analysis of the ligation of the NanoLuc reporter with GI-SmBiT (**A**) or with GI-SP (**B**). 50 μM of the ligation version NanoLuc (NanoLuc-NHV or NanoLuc-NAL), 500 μM of the ligation version peptide (GI-SmBiT or GI-SP), and 50 nM of the activated [G238V]BmAEP1 were mixed together in 200 mM phosphate buffer (pH 6.5). After incubation at 20 °C for the indicated time, 10 μl of the reaction mixture was removed and mixed with 40 μl of SDS-gel loading buffer. After boiling, 10 μl of the mixture was loaded onto a 15% SDS-gel and the gel was stained by Coomassie brilliant blue 250 after electrophoresis. (**C**) Saturation binding of the Luc-SP tracer with its receptor TACR1. To living HEK293T cells transiently overexpressing human TACR1, indicated concentrations of Luc-SP tracer with or without 1.0 μM of GI-SP were added (50 μl/well). After incubation at 22 °C for 1 h, the binding solution was removed and the cells were washed twice with ice-cold PBS. After addition of the diluted NanoLuc substrate (50 μl/well), bioluminescence was immediately measured on a SpectraMax iD3 plate reader. The measured bioluminescence data are expressed as mean ± SD (*n* = 3) and fitted to one-site binding model using the SigmaPlot10.0 software. **Inner panel**, SDS-PAGE analysis of the ligation product of NanoLuc-NAL with GI-SP. After 50 μM of NanoLuc-NAL was ligated with 500 μM of GI-SP by 50 nM of [G238V]BmAEP1 at 20 °C for 2 h, the ligation mixture was loaded onto a spin Ni^2+^ column. After washing by 30 mM imidazole, the bound fraction was eluted by 250 mM imidazole. Each fraction was loaded onto a 15% SDS-gel and the gel was stained by Coomassie brilliant blue 250 after electrophoresis.

Thereafter, we tested whether [G238V]BmAEP1 could ligate the NanoLuc reporter with neuropeptide substance P (SP), which has 11 amino acids (RPKPQQFFGLM) and an α-amidated C-terminus. C-terminal α-amidation is an important post-translational modification for many biologically active peptides. Although this post-translational modification remains challenging for recombinant expression, it is compatible with chemical peptide synthesis. Thus, enzymatic ligation of a synthetic peptide with the NanoLuc reporter provided a practical approach for preparation of bioluminescent tracers for these C-terminally α-amidated peptides. For this purpose, we introduced a nucleophilic motif (GIGG) into the N-terminus of human SP. SDS-PAGE analysis (Fig. 5B) showed that [G238V]BmAEP1 could efficiently catalyze the ligation of the two ligation version NanoLucs, especially NanoLuc-NAL, with the synthetic ligation version SP (GI-SP). After ligation, the excess amount of GI-SP was removed using a spin Ni^2+^ column, because it would drastically affect the receptor binding assay by functioning as a competitor. As analyzed by SDS-PAGE (Fig. 5C, inner panel), the eluted fraction from the Ni^2+^ column contained the expected ligation product Luc-SP (indicated by an asterisk) as well as a small amount of unreacted or hydrolyzed NanoLuc reporter. This eluted fraction was directly used for subsequent receptor binding assays because the small amount of unreacted or hydrolyzed NanoLuc had no significant influence on the binding of the Luc-SP tracer to its receptor.

We tested whether the Luc-SP tracer retains binding to its receptor TACR1, a G protein-coupled receptor, via a saturation binding assay. After Luc-SP was incubated with living HEK293T cells transiently overexpressing human TACR1, the measured bioluminescence increased in a hyperbolic manner as the tracer concentration increased (Fig. 5C). After competition using 1.0 μM of GI-SP, the measured bioluminescence decreased markedly. As calculated from the specific binding curve, the measured dissociation constant (K_d_) of Luc-SP with human TACR1 was 2.04 ± 0.14 nM (*n* = 3), and the measured maximal binding capacity (B_max_) was ∼2.5 × 10^6^ RLU/well, which was roughly equal to ∼5 × 10^4^ receptors/cell (the specific activity of Luc-SP was 5 × 10^5^ RLU/fmol, and each well contained ∼6 × 10^4^ cells). Thus, it seemed that the Luc-SP tracer could specifically bind to its receptor with high affinity, representing a novel non-radioactive tracer for ligand[receptor binding assays. In future studies, [G238V]BmAEP1 could be used to prepare NanoLuc-based binding tracers for other biologically active peptides.

### 3.7. Cyclization of the synthetic SFTI-NHV by [G238V]BmAEP1

To test whether [G238V]BmAEP1 could catalyze peptide cyclization, a synthetic precursor of sunflower trypsin inhibitor 1 (SFTI-1) was used as a model. For efficient ligation, the last Asp residue of the native SFTI-1 was replaced with an Asn residue and a His-Val dipeptide was introduced into the C-terminus of the synthetic precursor (Fig. 6A, inner panel). After chemical synthesis and purification, the precursor was subjected to oxidative refolding to form its intramolecular disulfide bond. After oxidization by 2,2’-dithiodipiridine in an arginine solution, the peak of the linear precursor disappeared and a new peak (indicated by an asterisk) with a slightly longer retention time appeared on HPLC (Fig. 6A). Mass spectrometry analysis demonstrated that the new peak displayed a molecular mass of 1766.8, consistent with the theoretical value (1767.1) of the refolded precursor. In Figure 6A, the peaks indicated by octothorpes are 2,2’-dithiodipiridine and its derived 2-pyridinethiol. After purification, the refolded SFTI-NHV displayed a symmetric elution peak and a slightly longer retention time, as analyzed by HPLC (Fig. S5).

**Fig. 6.**
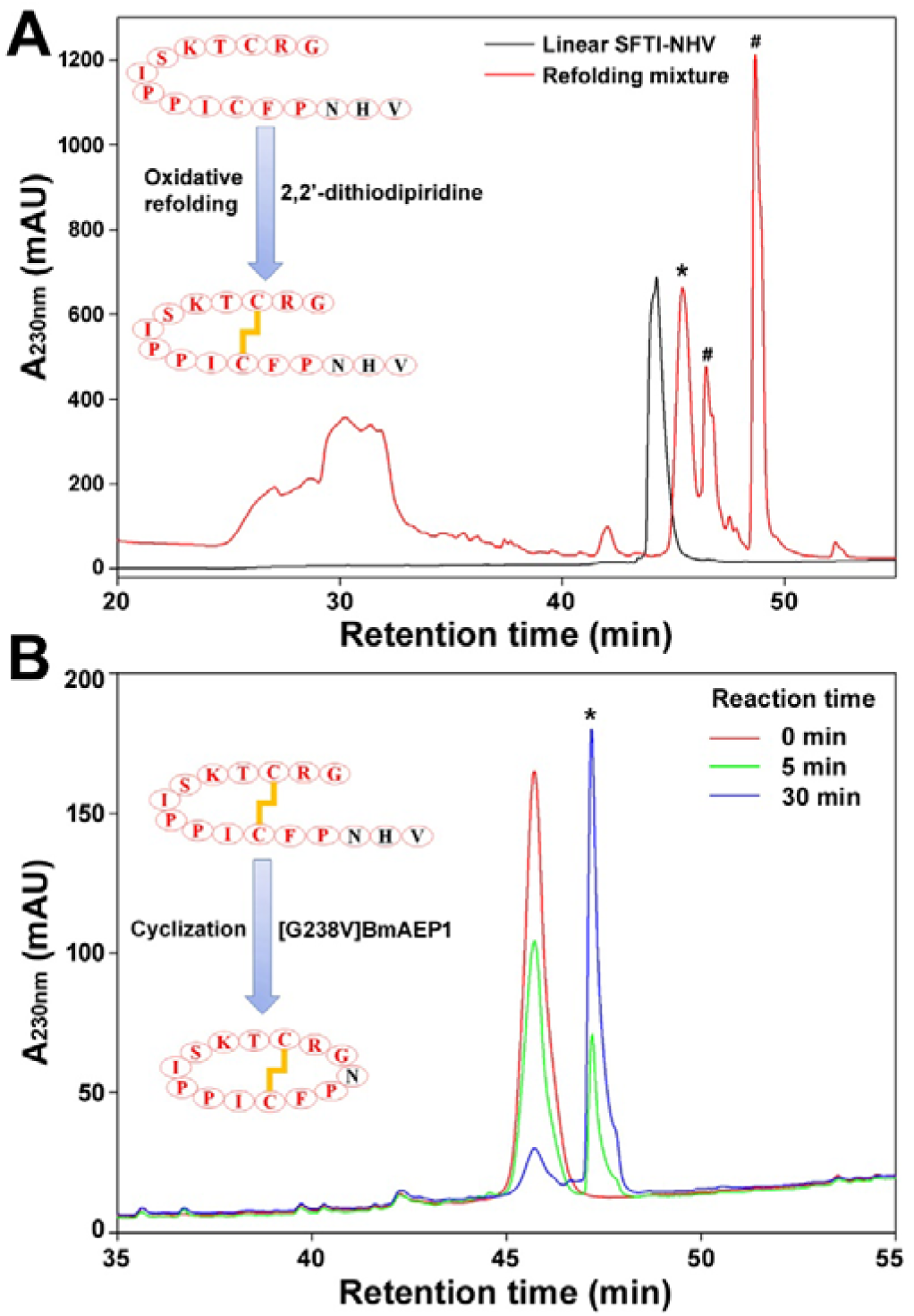
Cyclization of the synthetic SFTI-NHV by [G238V]BmAEP1. (**A**) HPLC analysis of the oxidative refolding of the linear SFTI-NHV. The synthetic linear peptide (∼100 μg) before or after refolding was loaded onto a C_18_ reverse-phase column and eluted by an acidic acetonitrile gradient. The peak of the refolded SFTI-NHV was indicated by an asterisk, those of 2,2’-dithiodipiridine and its derived 2-pyridinethiol were indicated by octothorpes. (**B**) HPLC analysis of the [G238V]BmAEP1-catalyzed cyclization of the folded SFTI-NHV. The refolded SFTI-NHV (∼20 μg) before or after cyclization was loaded onto a C_18_ reverse-phase column and eluted by an acidic acetonitrile gradient. The peak of the cyclic SFTI was indicated by an asterisk.

Subsequently, the refolded SFTI-NHV was cyclized by [G238V]BmAEP1. After 50 μM of the refolded SFTI-NHV was catalyzed by 50 nM of [G238V]BmAEP1, the peak of the precursor decreased quickly, and a new peak (indicated by an asterisk in the figure) with a longer retention time appeared and increased correspondingly, as analyzed using HPLC (Fig. 6B). Mass spectrometry analysis showed that the measured molecular mass (1512.6) of the new peak was consistent with the theoretical value (1512.9) of the cyclized SFTI. Thus, it seemed that [G238V]BmAEP1 could efficiently catalyze the intramolecular cyclization of synthetic peptides.

## Discussion

Using the newly developed NanoBiT-based peptide ligase activity assay, we demonstrated that AEP-type peptide ligase activity is widely present in bamboo species, a diverse group of evergreen perennial flowering plants in the subfamily of Bambusoideae. There are more than 1,400 bamboo species belonging to 115 genera [48]. Bamboos are distributed worldwide, from hot tropical regions to cool mountainous regions and high land cloud forests [48]. Thus, it seems that bamboos are a rich source for the identification of AEP-type peptide ligases in the future.

We identified an AEP-type peptide ligase, BmAEP1, from a popular horticultural bamboo species, *Bambusa multiplex*. According to its amino acid sequence, BmAEP1 belongs to the noncanonical AEP-type peptide ligases, such as McoAEP1 and McoAEP2 from *Momordica cochinchinensis* [37,38], because its residues at LAD1 and LAD2 are similar to those of AEP proteases, rather than those of the canonical AEP-type peptide ligases, such as butelase-1 and OaAEP1 (Fig. 1A). As a result, BmAEP1 displayed considerable hydrolysis activity, despite high peptide ligase activity. Introducing a G238V mutation at its LAD1 region markedly reduced its hydrolysis activity, but retained its ligase activity, confirming that LADs are responsible for the peptide ligase activity of AEPs [36,49]. In future studies, more AEPs might be converted to peptide ligases using this strategy [50].

AEPs are synthesized *in vivo* as inactive zymogens carrying a C-terminal cap domain (Fig. 1B). To remove the cap domain via cleavage of its peptide chain at the linker region, the catalytic core domain needs to exert its hydrolysis activity under acidic condition. Thus, AEP-type peptide ligases all have low hydrolysis activity, otherwise their zymogens cannot self-activate. As a result, it seems that natural selection cannot generate AEP-type peptide ligases that have completely lost the hydrolysis activity. For protein engineering, introduction of mutations at LADs can reduce AEP’s hydrolysis activity [50]; however, it might also impair self-activation of the mutant zymogens. As shown in the present study, the recombinant [G238V]BmAEP1 zymogen failed to self-activate, likely because of its low hydrolysis activity after the introduction of the mutation. Engineering of other AEPs might face the same problem.

To obtain soluble protein, intact AEP zymogens, rather than the catalytic core domain itself, are typically overexpressed in bacteria or other host cells, because the core domain alone is less stable and thus prone to forming inclusion bodies. To date, only one paper has reported the successful renaturation of the catalytic core domain of OaAEP1 from inclusion bodies [51]; however, whether this approach can be applied to other AEP-type peptide ligases is unknown. We have tested this approach using the core domain of butelase-1, but no ligase activity was detected after *in vitro* renaturation using the reported procedure. Thus, soluble overexpression of the intact zymogen is still a practical approach to prepare active AEP-type peptide ligases.

To activate the recombinant [G238V]BmAEP1 zymogen, herein, we developed a protease-assisted activation approach in which trypsin was used to cleave the recombinant zymogen and then was conveniently removed via an ion-exchange chromatography. Although there are many Lys and Arg residues in the core domain and cap domain, it seemed that trypsin specifically cleaved the peptide chain at the linker region, likely because of the flexible conformation of the linker region. This protease-assisted activation approach might be applied to other engineered AEP-type peptide ligases if they cannot self-activate because of low intrinsic hydrolysis activity. Besides trypsin, other endoproteinases might also be tested. If required, additional cleavage sites can be introduced into the linker region of the engineered zymogen for efficient *in vitro* cleavage of the engineered zymogens by a suitable endoproteinase.

## Acknowledgments

This work was supported by grants from the National Natural Science Foundation of China (31971193) and the Fundamental Research Funds for Platform of the International Centre for Bamboo and Rattan (1632021006).

## Notes

### Competing Interest Statement

The authors have declared no competing interest.

