## Supplemental Figures S1-S5 for "An efficient peptide ligase engineered from a bamboo asparaginyl endopeptidase"

**Contents:**

**Fig. S1.** The nucleotide and amino acid sequence of the ligation version NanoLuc reporters, NanoLuc-NHV and NanoLuc-NAL.

**Fig. S2.** The nucleotide and amino acid sequence of the BmAEP1 pre-zymogen deduced from the assembled transcriptomic sequencing data.

**Fig. S3.** The nucleotide and amino acid sequence of the N-terminally 6×His-tagged BmAEP1 zymogen overexpressed in *E. coli*.

**Fig. S4.** AEP-type peptide ligase activity detected in the crude extract of bamboo leaves.

**Fig. S5.** HPLC analysis of the unfolded SFTI-NHV and refolded SFTI-NHV.

**NanoLuc-NHV in pET vector**

```
1     ATG CAT CAC CAT CAC CAC CAT ACG GTC TTC ACA CTC GAA GAT TTC GTT GGG GAC TGG CGA CAG ACA GCC GGC TAC
      TAC GTA GTG GTA GTG GTG GTA TGC CAG AAG TGT GAG CTT CTA AAG CAA CCC CTG ACC GCT GTC TGT CGG CCG ATG
       M   H   H   H   H   H   H   T   V   F   T   L   E   D   F   V   G   D   W   R   Q   T   A   G   Y

76    AAC CTG GAC CAA GTC CTT GAA CAG GGA GGT GTG TCC AGT TTG TTT CAG AAT CTC GGG GTG TCC GTA ACT CCG ATC
      TTG GAC CTG GTT CAG GAA CTT GTC CCT CCA CAC AGG TCA AAC AAA GTC TTA GAG CCC CAC AGG CAT TGA GGC TAG
       N   L   D   Q   V   L   E   Q   G   G   V   S   S   L   F   Q   N   L   G   V   S   V   T   P   I

151   CAA AGG ATT GTC CTG AGC GGT GAA AAT GGG CTG AAG ATC GAC ATC CAT GTC ATC ATC CCG TAT GAA GGT CTG AGC
      GTT TCC TAA CAG GAC TCG CCA CTT TTA CCC GAC TTC TAG CTG TAG GTA CAG TAG TAG GGC ATA CTT CCA GAC TCG
       Q   R   I   V   L   S   G   E   N   G   L   K   I   D   I   H   V   I   I   P   Y   E   G   L   S

226   GGC GAC CAA ATG GGC CAG ATC GAA AAA ATT TTT AAG GTG GTG TAC CCT GTG GAT GAT CAT CAC TTT AAG GTG ATC
      CCG CTG GTT TAC CCG GTC TAG CTT TTT TAA AAA TTC CAC CAC ATG GGA CAC CTA CTA GTA GTG AAA TTC CAC TAG
       G   D   Q   M   G   Q   I   E   K   I   F   K   V   V   Y   P   V   D   D   H   H   F   K   V   I

301   CTG CAC TAT GGC ACA CTG GTA ATC GAC GGG GTT ACG CCG AAC ATG ATC GAC TAT TTC GGA CGG CCG TAT GAA GGC
      GAC GTG ATA CCG TGT GAC CAT TAG CTG CCC CAA TGC GGC TTG TAC TAG CTG ATA AAG CCT GCC GGC ATA CTT CCG
       L   H   Y   G   T   L   V   I   D   G   V   T   P   N   M   I   D   Y   F   G   R   P   Y   E   G

376   ATC GCC GTG TTC GAC GGC AAA AAG ATC ACT GTA ACA GGG ACC CTG TGG AAC GGC AAC AAA ATT ATC GAC GAG CGC
      TAG CGG CAC AAG CTG CCG TTT TTC TAG TGA CAT TGT CCC TGG GAC ACC TTG CCG TTG TTT TAA TAG CTG CTC GCG
       I   A   V   F   D   G   K   K   I   T   V   T   G   T   L   W   N   G   N   K   I   I   D   E   R

451   CTG ATC AAC CCC GAC GGC TCC CTG CTG TTC CGA GTA ACC ATC CAA GGA GTG ACC GGC TGG CGG CTG TGC GAA CGC
      GAC TAG TTG GGG CTG CCG AGG GAC GAC AAG GCT CAT TGG TAG GTT CCT CAC TGG CCG ACC GCC GAC ACG CTT GCG
       L   I   N   P   D   G   S   L   L   F   R   V   T   I   Q   G   V   T   G   W   R   L   C   E   R

526   ATT CTG GCG GGT GGC GGT GGC CAT ATG GGT GGC AGC GGC GGT GGT AGT GGT GGC GGT GGC GGC AAC CAT GTG TAA
      TAA GAC CGC CCA CCG CCA CCG GTA TAC CCA CCG TCG CCG CCA CCA TCA CCA CCG CCA CCG CCG TTG GTA CAC ATT
       I   L   A   G   G   G   G   H   M   G   G   S   G   G   G   S   G   G   G   G   G   N   H   V   *
```

**NanoLuc-NAL in pET vector**

```
1     ATG CAT CAC CAT CAC CAC CAT ACG GTC TTC ACA CTC GAA GAT TTC GTT GGG GAC TGG CGA CAG ACA GCC GGC TAC
      TAC GTA GTG GTA GTG GTG GTA TGC CAG AAG TGT GAG CTT CTA AAG CAA CCC CTG ACC GCT GTC TGT CGG CCG ATG
       M   H   H   H   H   H   H   T   V   F   T   L   E   D   F   V   G   D   W   R   Q   T   A   G   Y

76    AAC CTG GAC CAA GTC CTT GAA CAG GGA GGT GTG TCC AGT TTG TTT CAG AAT CTC GGG GTG TCC GTA ACT CCG ATC
      TTG GAC CTG GTT CAG GAA CTT GTC CCT CCA CAC AGG TCA AAC AAA GTC TTA GAG CCC CAC AGG CAT TGA GGC TAG
       N   L   D   Q   V   L   E   Q   G   G   V   S   S   L   F   Q   N   L   G   V   S   V   T   P   I

151   CAA AGG ATT GTC CTG AGC GGT GAA AAT GGG CTG AAG ATC GAC ATC CAT GTC ATC ATC CCG TAT GAA GGT CTG AGC
      GTT TCC TAA CAG GAC TCG CCA CTT TTA CCC GAC TTC TAG CTG TAG GTA CAG TAG TAG GGC ATA CTT CCA GAC TCG
       Q   R   I   V   L   S   G   E   N   G   L   K   I   D   I   H   V   I   I   P   Y   E   G   L   S

226   GGC GAC CAA ATG GGC CAG ATC GAA AAA ATT TTT AAG GTG GTG TAC CCT GTG GAT GAT CAT CAC TTT AAG GTG ATC
      CCG CTG GTT TAC CCG GTC TAG CTT TTT TAA AAA TTC CAC CAC ATG GGA CAC CTA CTA GTA GTG AAA TTC CAC TAG
       G   D   Q   M   G   Q   I   E   K   I   F   K   V   V   Y   P   V   D   D   H   H   F   K   V   I

301   CTG CAC TAT GGC ACA CTG GTA ATC GAC GGG GTT ACG CCG AAC ATG ATC GAC TAT TTC GGA CGG CCG TAT GAA GGC
      GAC GTG ATA CCG TGT GAC CAT TAG CTG CCC CAA TGC GGC TTG TAC TAG CTG ATA AAG CCT GCC GGC ATA CTT CCG
       L   H   Y   G   T   L   V   I   D   G   V   T   P   N   M   I   D   Y   F   G   R   P   Y   E   G

376   ATC GCC GTG TTC GAC GGC AAA AAG ATC ACT GTA ACA GGG ACC CTG TGG AAC GGC AAC AAA ATT ATC GAC GAG CGC
      TAG CGG CAC AAG CTG CCG TTT TTC TAG TGA CAT TGT CCC TGG GAC ACC TTG CCG TTG TTT TAA TAG CTG CTC GCG
       I   A   V   F   D   G   K   K   I   T   V   T   G   T   L   W   N   G   N   K   I   I   D   E   R

451   CTG ATC AAC CCC GAC GGC TCC CTG CTG TTC CGA GTA ACC ATC CAA GGA GTG ACC GGC TGG CGG CTG TGC GAA CGC
      GAC TAG TTG GGG CTG CCG AGG GAC GAC AAG GCT CAT TGG TAG GTT CCT CAC TGG CCG ACC GCC GAC ACG CTT GCG
       L   I   N   P   D   G   S   L   L   F   R   V   T   I   Q   G   V   T   G   W   R   L   C   E   R

526   ATT CTG GCG GGT GGC GGT GGC CAT ATG GGT GGC AGC GGC GGT GGT AGT GGT GGC GGT GGC GGC AAC GCT CTG TAA
      TAA GAC CGC CCA CCG CCA CCG GTA TAC CCA CCG TCG CCG CCA CCA TCA CCA CCG CCA CCG CCG TTG CGA GAC ATT
       I   L   A   G   G   G   G   H   M   G   G   S   G   G   G   S   G   G   G   G   G   N   A   L   *
```

**Fig. S1.** The nucleotide and amino acid sequence of the ligation version NanoLuc reporters, NanoLuc-NHV and NanoLuc-NAL. The amino acid sequence of NanoLuc is shown in blue, the recognition motif of the AEP-type peptide ligase is shaded.

```
1     ATG GCG TCT ATC CGC CTT CTT CCC CTC GCG CTG CTG CTC TCC GCG CTC GCC GTC GTG CAG GCC CGG TGG CCG ACC
      TAC CGC AGA TAG GCG GAA GAA GGG GAG CGC GAC GAC GAG AGG CGC GAG CGG CAG CAC GTC CGG GCC ACC GGC TGG
1      M   A   S   I   R   L   L   P   L   A   L   L   L   S   A   L   A   V   V   Q   A   R   W   P   T

76    ATC CGG CTG CCG TCG GAG CGC GCC GCG GAC GAG GCG GAG GAG GAG GAG GAC GAC TCC GTC GGG ACC AGG TGG GCC
      TAG GCC GAC GGC AGC CTC GCG CGG CGC CTG CTC CGC CTC CTC CTC CTC CTG CTG AGG CAG CCC TGG TCC ACC CGG
26     I   R   L   P   S   E   R   A   A   D   E   A   E   E   E   E   D   D   S   V   G   T   R   W   A

151   GTC CTC GTC GCC GGC TCC AAC GGG TAC TAC AAC TAC CGC CAC CAG GCG GAT ATC TGC CAC GCC TAC CAG ATC ATG
      CAG GAG CAG CGG CCG AGG TTG CCC ATG ATG TTG ATG GCG GTG GTC CGC CTA TAG ACG GTG CGG ATG GTC TAG TAC
51     V   L   V   A   G   S   N   G   Y   Y   N   Y   R   H   Q   A   D   I   C   H   A   Y   Q   I   M

226   AAG AAG GGC GGG CTC AAG GAC GAG AAC ATC ATT GTC TTC ATG TAC GAT GAC ATC GCG CAC AGC CCG GAG AAT CCG
      TTC TTC CCG CCC GAG TTC CTG CTC TTG TAG TAA CAG AAG TAC ATG CTA CTG TAG CGC GTG TCG GGC CTC TTA GGC
76     K   K   G   G   L   K   D   E   N   I   I   V   F   M   Y   D   D   I   A   H   S   P   E   N   P

301   AGG CCT GGT GTC ATC ATC AAC CAT CCC CAG GGT GGC GAT GTC TAT GCT GGG GTC CCA AAG GAT TAC ACC GGG AAG
      TCC GGA CCA CAG TAG TAG TTG GTA GGG GTC CCA CCG CTA CAG ATA CGA CCC CAG GGT TTC CTA ATG TGG CCC TTC
101    R   P   G   V   I   I   N   H   P   Q   G   G   D   V   Y   A   G   V   P   K   D   Y   T   G   K

376   GAG GTT AGT GTC AAT AAC TTC TTC GCT GTT CTG CTC GGT AAT AAA ACC GCT GTC AGT GGT GGG AGC GGC AAA GTC
      CTC CAA TCA CAG TTA TTG AAG AAG CGA CAA GAC GAG CCA TTA TTT TGG CGA CAG TCA CCA CCC TCG CCG TTT CAG
126    E   V   S   V   N   N   F   F   A   V   L   L   G   N   K   T   A   V   S   G   G   S   G   K   V

451   GTG GAC AGT GGC CCC AAC GAT CAT ATT TTC ATT TTC TAC AGT GAC CAT GGG GGT CCT GGT GTC CTT GGG ATG CCT
      CAC CTG TCA CCG GGG TTG CTA GTA TAA AAG TAA AAG ATG TCA CTG GTA CCC CCA GGA CCA CAG GAA CCC TAC GGA
151    V   D   S   G   P   N   D   H   I   F   I   F   Y   S   D   H   G   G   P   G   V   L   G   M   P

526   ACC TAC CCA TAC CTC TAC GGT GAC GAC CTC GTA GAT ATT CTG AAG AAG AAG CAC GCT GCT GGA ACC TAC AAA AGC
      TGG ATG GGT ATG GAG ATG CCA CTG CTG GAG CAT CTA TAA GAC TTC TTC TTC GTG CGA CGA CCT TGG ATG TTT TCG
176    T   Y   P   Y   L   Y   G   D   D   L   V   D   I   L   K   K   K   H   A   A   G   T   Y   K   S

601   CTG GTC TTT TAC CTT GAA GCC TGC GAA TCC GGG AGC ATC TTT GAG GGC CTT CTG CCG AAT GAC ATC GAC GTC TAC
      GAC CAG AAA ATG GAA CTT CGG ACG CTT AGG CCC TCG TAG AAA CTC CCG GAA GAC GGC TTA CTG TAG CTG CAG ATG
201    L   V   F   Y   L   E   A   C   E   S   G   S   I   F   E   G   L   L   P   N   D   I   D   V   Y

676   GCG ACC ACC GCA TCA AAC GCA GAG GAG AGC AGC TGG GGG ACG TAC TGC CCC GGC GAG TAC CCG AGC CCT CCT CCG
      CGC TGG TGG CGT AGT TTG CGT CTC CTC TCG TCG ACC CCC TGC ATG ACG GGG CCG CTC ATG GGC TCG GGA GGA GGC
226    A   T   T   A   S   N   A   E   E   S   S   W   G   T   Y   C   P   G   E   Y   P   S   P   P   P

751   GAG TAC GAC ACC TGC TTG GGG GAC TTG TAC AGC GTT TCT TGG ATG GAA GAC AGC GAT GTC CAC AAC CTG CGA ACA
      CTC ATG CTG TGG ACG AAC CCC CTG AAC ATG TCG CAA AGA ACC TAC CTT CTG TCG CTA CAG GTG TTG GAC GCT TGT
251    E   Y   D   T   C   L   G   D   L   Y   S   V   S   W   M   E   D   S   D   V   H   N   L   R   T

826   GAA TCT CTC AAG CAG CAG TAC AAG CTG GTC AAG GAC AGG ACA TCG GTT CAG AAC ACA TAC AGC TAT GGT TCC CAT
      CTT AGA GAG TTC GTC GTC ATG TTC GAC CAG TTC CTG TCC TGT AGC CAA GTC TTG TGT ATG TCG ATA CCA AGG GTA
276    E   S   L   K   Q   Q   Y   K   L   V   K   D   R   T   S   V   Q   N   T   Y   S   Y   G   S   H

901   GTG ATG CAA TAC GGT TCT TTG GAC CTG AAT GTT CAA CAT CTG TTC TTG TAC ATC GGA TCA AAT CCT GCT AAC GAG
      CAC TAC GTT ATG CCA AGA AAC CTG GAC TTA CAA GTT GTA GAC AAG AAC ATG TAG CCT AGT TTA GGA CGA TTG CTC
301    V   M   Q   Y   G   S   L   D   L   N   V   Q   H   L   F   L   Y   I   G   S   N   P   A   N   E

976   AAC GCC ACG TTT GTG GAA GAC AAC TCA TTG CCG TCG TTC TCA AGA GCT GTT AAT CAG AGA GAT GCT GAT CTT GTT
      TTG CGG TGC AAA CAC CTT CTG TTG AGT AAC GGC AGC AAG AGT TCT CGA CAA TTA GTC TCT CTA CGA CTA GAA CAA
326    N   A   T   F   V   E   D   N   S   L   P   S   F   S   R   A   V   N   Q   R   D   A   D   L   V

1051  TAC TTC TGG CAG AAG TAC CGG AAA TTG GCC GAG GGT TCC CCT GAG AAA AAC AAT GCT CGG AAG CAA TTG CTT GAA
      ATG AAG ACC GTC TTC ATG GCC TTT AAC CGG CTC CCA AGG GGA CTC TTT TTG TTA CGA GCC TTC GTT AAC GAA CTT
351    Y   F   W   Q   K   Y   R   K   L   A   E   G   S   P   E   K   N   N   A   R   K   Q   L   L   E

1126  GTG ATG GCC CAT AGA TCT CAT GTT GAC AAC AGT GTC AAG CTG ATC GGA AAC CTT CTG TTT GGC TCT GAG GAT GGT
      CAC TAC CGG GTA TCT AGA GTA CAA CTG TTG TCA CAG TTC GAC TAG CCT TTG GAA GAC AAA CCG AGA CTC CTA CCA
376    V   M   A   H   R   S   H   V   D   N   S   V   K   L   I   G   N   L   L   F   G   S   E   D   G

1201  CCA AGG GTT CTG AAG ACT GTT CGT GCA GCT GGT GAA CCT CTG GTT AAT GAC TGG ACC TGC CTC AAG TCT ATG GTG
      GGT TCC CAA GAC TTC TGA CAA GCA CGT CGA CCA CTT GGA GAC CAA TTA CTG ACC TGG ACG GAG TTC AGA TAC CAC
401    P   R   V   L   K   T   V   R   A   A   G   E   P   L   V   N   D   W   T   C   L   K   S   M   V

1276  CGT GCT TTT GAA GCG CAA TGT GGC TCG TTG GCG CAG TAT GGA ATG AAG CAC ATG CGG TCC TTT GCA AAC ATC TGC
```

GCA CGA AAA CTT CGC GTT ACA CCG AGC AAC CGC GTC ATA CCT TAC TTC GTG TAC GCC AGG AAA CGT TTG TAG ACG
426 R A F E A Q C G S L A Q Y G M K H M R S F A N I C

1351 AAT GCT GGC ATC CGT GTT GAA GCG ATG GCA AAG GTT GCC GCG CAG GCT TGC ACG AGC ATT CCC TCC AAC CCC TGG
TTA CGA CCG TAG GCA CAA CTT CGC TAC CGT TTC CAA CGG CGC GTC CGA ACG TGC TCG TAA GGG AGG TTG GGG ACC
451 N A G I R V E A M A K V A A Q A C T S I P S N P W

1426 AGT TCC ATC CAC AAG GGT TTT AGT GCT TGA
TCA AGG TAG GTG TTC CCA AAA TCA CGA ACT
476 S S I H K G F S A *

**Fig. S2.** The nucleotide and amino acid sequence of the BmAEP1 pre-zymogen deduced from the assembled transcriptomic sequencing data. The predicted signal peptide (residues 1-21) is shaded. The positions of the cDNA cloning primers are underlined. For cDNA cloning, three transformed *E. coli* colonies were sequenced, one cDNA bears one missense mutation (tat to ttg/Y297L) and two synonymous mutations (t417c; c486t), the other two cDNAs are identical and bear two missense mutations (cac to tac/H95Y; tat to ttg/Y297L) and nine synonymous mutations (c246t; t258c; c279t; g282t; t306c; t309c; t333g; t417c; c486t) compared to the transcriptomic sequencing-derived BmAEP1 pre-zymogen.

```
                                  Nde I
1     ATG CAT CAC CAT CAC CAC CAT ATG CGG TGG CCG ACC ATC CGG CTG CCG TCG GAG CGC GCC GCG GAC GAG GCG GAG
      TAC GTA GTG GTA GTG GTG GTA TAC GCC ACC GGC TGG TAG GCC GAC GGC AGC CTC GCG CGG CGC CTG CTC CGC CTC
       M   H   H   H   H   H   H   M   R   W   P   T   I   R   L   P   S   E   R   A   A   D   E   A   E

76    GAG GAG GAG GAC GAC TCC GTC GGG ACC CGT TGG GCC GTC CTC GTC GCC GGC TCC AAC GGG TAC TAC AAC TAC CGC
      CTC CTC CTC CTG CTG AGG CAG CCC TGG GCA ACC CGG CAG GAG CAG CGG CCG AGG TTG CCC ATG ATG TTG ATG GCG
       E   E   E   D   D   S   V   G   T   R   W   A   V   L   V   A   G   S   N   G   Y   Y   N   Y   R

151   CAC CAG GCG GAT ATC TGC CAC GCC TAC CAG ATC ATG AAG AAG GGC GGG CTC AAG GAC GAG AAC ATC ATT GTC TTC
      GTG GTC CGC CTA TAG ACG GTG CGG ATG GTC TAG TAC TTC TTC CCG CCC GAG TTC CTG CTC TTG TAG TAA CAG AAG
       H   Q   A   D   I   C   H   A   Y   Q   I   M   K   K   G   G   L   K   D   E   N   I   I   V   F

226   ATG TAC GAT GAC ATC GCG CAC AGC CCG GAG AAT CCG CGC CCT GGT GTC ATC ATC AAC CAT CCC CAG GGT GGC GAT
      TAC ATG CTA CTG TAG CGC GTG TCG GGC CTC TTA GGC GCG GGA CCA CAG TAG TAG TTG GTA GGG GTC CCA CCG CTA
       M   Y   D   D   I   A   H   S   P   E   N   P   R   P   G   V   I   I   N   H   P   Q   G   G   D

301   GTC TAT GCT GGG GTC CCA AAG GAT TAC ACC GGG AAG GAG GTT AGT GTC AAT AAC TTC TTC GCT GTT CTG CTC GGT
      CAG ATA CGA CCC CAG GGT TTC CTA ATG TGG CCC TTC CTC CAA TCA CAG TTA TTG AAG AAG CGA CAA GAC GAG CCA
       V   Y   A   G   V   P   K   D   Y   T   G   K   E   V   S   V   N   N   F   F   A   V   L   L   G

376   AAT AAA ACC GCT GTC AGT GGT GGG AGC GGC AAA GTC GTG GAC AGT GGC CCC AAC GAT CAT ATT TTC ATT TTC TAC
      TTA TTT TGG CGA CAG TCA CCA CCC TCG CCG TTT CAG CAC CTG TCA CCG GGG TTG CTA GTA TAA AAG TAA AAG ATG
       N   K   T   A   V   S   G   G   S   G   K   V   V   D   S   G   P   N   D   H   I   F   I   F   Y

451   AGT GAC CAT GGG GGT CCT GGT GTC CTT GGG ATG CCT ACC TAC CCA TAC CTC TAC GGT GAC GAC CTC GTA GAT ATT
      TCA CTG GTA CCC CCA GGA CCA CAG GAA CCC TAC GGA TGG ATG GGT ATG GAG ATG CCA CTG CTG GAG CAT CTA TAA
       S   D   H   G   G   P   G   V   L   G   M   P   T   Y   P   Y   L   Y   G   D   D   L   V   D   I

526   CTG AAG AAG AAG CAC GCT GCT GGA ACC TAC AAA AGC CTG GTC TTT TAC CTT GAA GCC TGC GAA TCC GGG AGC ATC
      GAC TTC TTC TTC GTG CGA CGA CCT TGG ATG TTT TCG GAC CAG AAA ATG GAA CTT CGG ACG CTT AGG CCC TCG TAG
       L   K   K   K   H   A   A   G   T   Y   K   S   L   V   F   Y   L   E   A   C   E   S   G   S   I

601   TTT GAG GGC CTT CTG CCG AAT GAC ATC GAC GTC TAC GCG ACC ACC GCA TCA AAC GCA GAG GAG AGC AGC TGG GGG
      AAA CTC CCG GAA GAC GGC TTA CTG TAG CTG CAG ATG CGC TGG TGG CGT AGT TTG CGT CTC CTC TCG TCG ACC CCC
       F   E   G   L   L   P   N   D   I   D   V   Y   A   T   T   A   S   N   A   E   E   S   S   W   G

676   ACG TAC TGC CCC GGC GAG TAC CCG AGC CCT CCT CCG GAG TAC GAC ACC TGC TTG GGG GAC TTG TAC AGC GTT TCT
      TGC ATG ACG GGG CCG CTC ATG GGC TCG GGA GGA GGC CTC ATG CTG TGG ACG AAC CCC CTG AAC ATG TCG CAA AGA
       T   Y   C   P   G   E   Y   P   S   P   P   P   E   Y   D   T   C   L   G   D   L   Y   S   V   S

751   TGG ATG GAA GAC AGC GAT GTC CAC AAC CTG CGT ACA GAA TCT CTC AAG CAG CAG TAC AAG CTG GTC AAG GAC CGT
      ACC TAC CTT CTG TCG CTA CAG GTG TTG GAC GCA TGT CTT AGA GAG TTC GTC GTC ATG TTC GAC CAG TTC CTG GCA
       W   M   E   D   S   D   V   H   N   L   R   T   E   S   L   K   Q   Q   Y   K   L   V   K   D   R

826   ACA TCG GTT CAG AAC ACA TAC AGC TAT GGT TCC CAT GTG ATG CAA TAC GGT TCT TTG GAC CTG AAT GTT CAA CAT
      TGT AGC CAA GTC TTG TGT ATG TCG ATA CCA AGG GTA CAC TAC GTT ATG CCA AGA AAC CTG GAC TTA CAA GTT GTA
       T   S   V   Q   N   T   Y   S   Y   G   S   H   V   M   Q   Y   G   S   L   D   L   N   V   Q   H

901   CTG TTC TTG TAC ATC GGA TCA AAT CCT GCT AAC GAG AAC GCC ACG TTT GTG GAA GAC AAC TCA TTG CCG TCG TTC
      GAC AAG AAC ATG TAG CCT AGT TTA GGA CGA TTG CTC TTG CGG TGC AAA CAC CTT CTG TTG AGT AAC GGC AGC AAG
       L   F   L   Y   I   G   S   N   P   A   N   E   N   A   T   F   V   E   D   N   S   L   P   S   F

976   TCA AGA GCT GTT AAT CAG AGA GAT GCT GAT CTT GTT TAC TTC TGG CAG AAG TAC CGG AAA TTG GCC GAG GGT TCC
      AGT TCT CGA CAA TTA GTC TCT CTA CGA CTA GAA CAA ATG AAG ACC GTC TTC ATG GCC TTT AAC CGG CTC CCA AGG
       S   R   A   V   N   Q   R   D   A   D   L   V   Y   F   W   Q   K   Y   R   K   L   A   E   G   S

1051  CCT GAG AAA AAC AAT GCT CGG AAG CAA TTG CTT GAA GTG ATG GCC CAT AGA TCT CAT GTT GAC AAC AGT GTC AAG
      GGA CTC TTT TTG TTA CGA GCC TTC GTT AAC GAA CTT CAC TAC CGG GTA TCT AGA GTA CAA CTG TTG TCA CAG TTC
       P   E   K   N   N   A   R   K   Q   L   L   E   V   M   A   H   R   S   H   V   D   N   S   V   K

1126  CTG ATC GGA AAC CTT CTG TTT GGC TCT GAG GAT GGT CCA CGC GTT CTG AAG ACT GTT CGT GCA GCT GGT GAA CCT
      GAC TAG CCT TTG GAA GAC AAA CCG AGA CTC CTA CCA GGT GCG CAA GAC TTC TGA CAA GCA CGT CGA CCA CTT GGA
       L   I   G   N   L   L   F   G   S   E   D   G   P   R   V   L   K   T   V   R   A   A   G   E   P

1201  CTG GTT AAT GAC TGG ACC TGC CTC AAG TCT ATG GTG CGT GCT TTT GAA GCG CAA TGT GGC TCG TTG GCG CAG TAT
      GAC CAA TTA CTG ACC TGG ACG GAG TTC AGA TAC CAC GCA CGA AAA CTT CGC GTT ACA CCG AGC AAC CGC GTC ATA
       L   V   N   D   W   T   C   L   K   S   M   V   R   A   F   E   A   Q   C   G   S   L   A   Q   Y

1276  GGA ATG AAG CAC ATG CGG TCC TTT GCA AAC ATC TGC AAT GCT GGC ATC CGT GTT GAA GCG ATG GCA AAG GTT GCC
      CCT TAC TTC GTG TAC GCC AGG AAA CGT TTG TAG ACG TTA CGA CCG TAG GCA CAA CTT CGC TAC CGT TTC CAA CGG
       G   M   K   H   M   R   S   F   A   N   I   C   N   A   G   I   R   V   E   A   M   A   K   V   A

                                                                                          EcoRI
1351  GCG CAG GCT TGC ACG AGC ATT CCC TCC AAC CCC TGG AGT TCC ATC CAC AAG GGT TTT AGT GCT TAA GAA TTC
      CGC GTC CGA ACG TGC TCG TAA GGG AGG TTG GGG ACC TCA AGG TAG GTG TTC CCA AAA TCA CGA ATT CTT AAG
       A   Q   A   C   T   S   I   P   S   N   P   W   S   S   I   H   K   G   F   S   A   *
```

**Fig. S3.** The nucleotide and amino acid sequence of the N-terminally 6×His-tagged BmAEP1 zymogen overexpressed in *E. coli*. The restriction enzyme cleavage sites for cloning of the synthetic gene to the pET expression vector are shaded. The amino acids in LAD1 and LAD2 are shown in red and shaded.

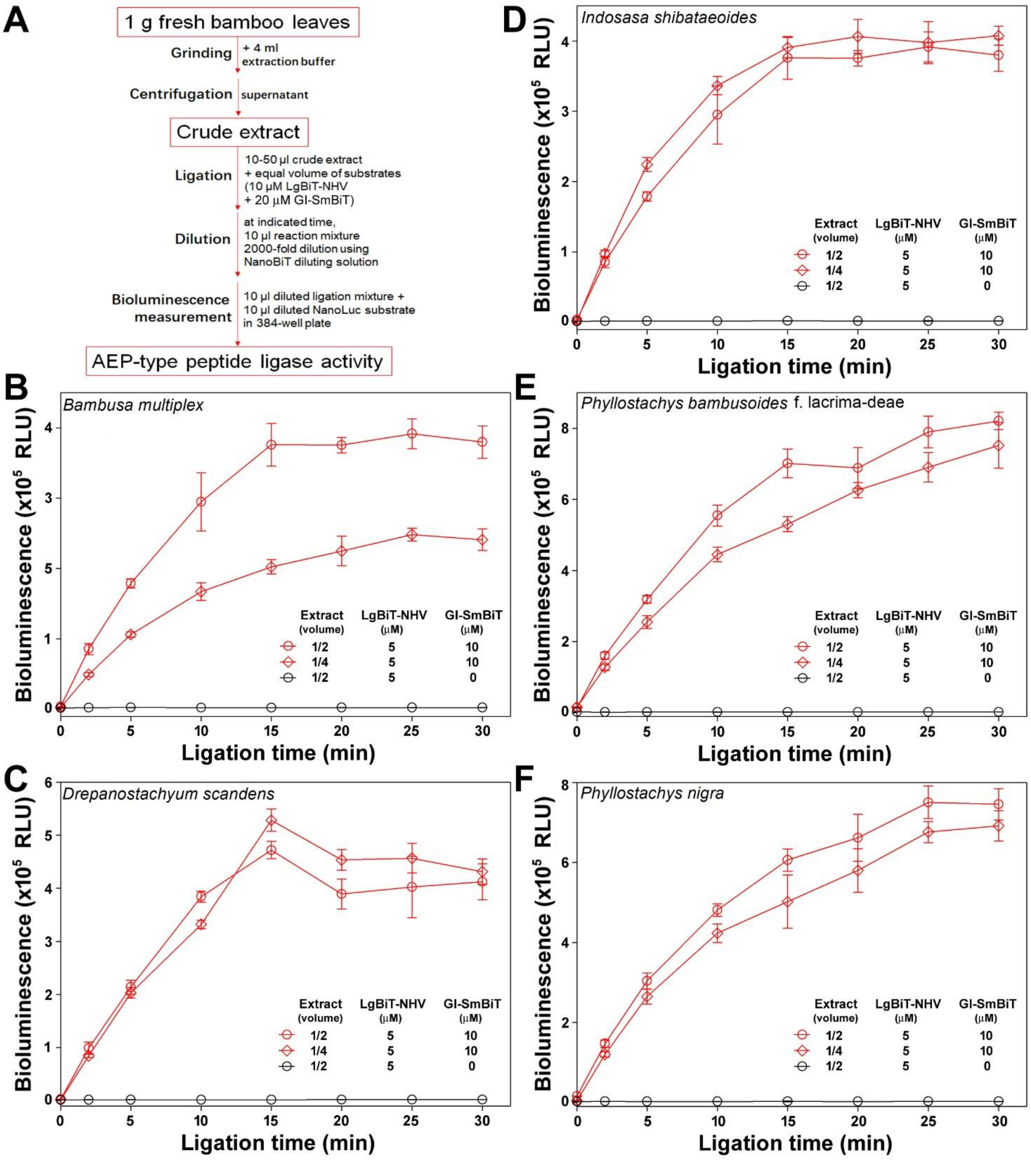


**Fig. S4.** AEP-type peptide ligase activity detected in the crude extract of bamboo leaves. (**A**) Procedure for the NanoBiT-based peptide ligase activity assay. (**B–F**) AEP-type peptide ligase activity detected in the crude extract of some bamboo leaves.

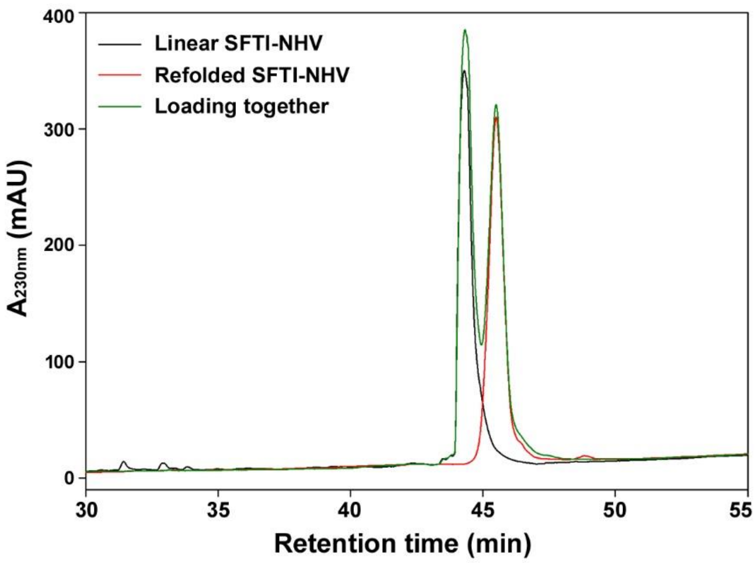


**Fig. S5.** HPLC analysis of the linear SFTI-NHV and the refolded SFTI-NHV.
